# RESPAN: A Deep Learning Pipeline for Accurate and Automated Restoration, Segmentation, and Quantification of Dendritic Spines

**DOI:** 10.1101/2024.06.06.597812

**Authors:** Sergio B. Garcia, Alexa P. Schlotter, Daniela Pereira, Aleksandra J. Recupero, Franck Polleux, Luke A. Hammond

## Abstract

Quantification of dendritic spines is essential for studying synaptic connectivity, yet most current approaches require manual adjustments or the combination of multiple software tools for optimal results. Here, we present <u>R</u>estoration <u>E</u>nhanced <u>SP</u>ine <u>A</u>nd <u>N</u>euron Analysis (RESPAN), an open-source pipeline integrating state-of-the-art deep learning for image restoration, segmentation, and analysis in an easily deployable, user-friendly interface. Leveraging content-aware restoration to enhance signal, contrast, and isotropic resolution further enhances RESPAN’s robust detection of spines, dendritic branches, and soma across a wide variety of samples, including challenging datasets such as those from live imaging and in vivo 2-photon microscopy with limited signal. Extensive validation against expert annotations and comparison with other software demonstrates RESPAN’s superior accuracy and reproducibility across multiple imaging modalities. RESPAN offers significant improvements in usability over currently available approaches, streamlining and democratizing access to a combination of advanced capabilities through an accessible resource for the neuroscience community.

**MOTIVATION:** Accurate and unbiased reconstructions of neuronal morphology and quantification of dendritic spines are widely used in neuroscience but remain a significant challenge for efficient large-scale analysis. Current methods rely heavily on parameter optimization between images and manual annotation, introducing bias and creating bottlenecks that limit large-scale studies. Additionally, existing automated tools often require complex workflows across multiple software platforms and lack integrated validation capabilities. We developed RESPAN to address these limitations by providing a comprehensive, automated pipeline that combines state-of-the-art deep learning approaches for image restoration, model training, image segmentation and analysis within one user-friendly graphic interface. This enables rapid, unbiased analysis of dendritic spine morphology across diverse imaging modalities while maintaining high accuracy and reproducibility.

## INTRODUCTION

Dendritic morphology and synaptic input connectivity are among the cardinal features defining neuronal subtypes and their functional properties ^1,2^. In the mammalian neocortex, pyramidal neurons (PNs) receive the majority (>90%) of their excitatory synaptic inputs on dendritic spines – small protrusions first described by Santiago Ramón y Cajal ^3,4^. Analysis of dendritic spine density and distribution has been used for decades as a critical anatomical proxy for assessing excitatory input across different PN subtypes during development and in adult states, both in physiological and pathological contexts, including neurodevelopmental disorders and neurodegeneration ^5–7^. Furthermore, the size of spine heads correlates linearly with postsynaptic AMPA receptor content, making it a key indicator of synaptic maturation during development and synaptic strength in adults ^8^.

While recent deep learning (DL) approaches have improved the accuracy of dendritic spine mapping capabilities ^9–11^, their utility remains constrained by limited functionality and analysis readouts. Existing methods like Vaa3D, SpineTool, and Imaris ^1,12–16^, Imaris (RRID:SCR_007370)) lack robustness and scalability across datasets and are highly sensitive to changes in signal and image quality. Consequently, full automation is rarely achieved, as each image often requires extensive manual intervention for parameter optimization and post-processing corrections. This reliance on manual adjustments limits efficiency and introduces human error and bias. While these tools can be combined with DL image restoration and segmentation tools, doing so requires significant technical expertise and creates complex, multi-step workflows that limit throughput and broader adoption.

In recent decades, two key developments have made transformative advances in dendritic spine imaging. First, the introduction of bright fluorescent proteins combined with improved expression methods, including in utero electroporation (IUE), viral delivery, and transgenic mouse models, has enabled sparse, cell-type specific labeling ^17–19^. Second, improvements in single and multi-photon confocal microscopy have dramatically enhanced spatial and temporal resolution for both in vitro and in vivo imaging ^20,21^. However, accurate quantification of dendritic morphology, spine density, 3D distribution, and spine head size remains challenging due to reliance on labor-intensive manual or semi-automated reconstructions, which introduce subjectivity and constitute significant bottlenecks for large-scale connectivity studies.

To address these challenges, we developed <u>R</u>estoration <u>E</u>nhanced <u>SP</u>ine <u>A</u>nd <u>N</u>euron Analysis (RESPAN), an integrated solution for automated neuron and dendritic spine analysis. RESPAN uniquely combines state-of-the-art deep learning approaches for content-aware image restoration, enhanced axial resolution, and robust 3D segmentation within a single, user-friendly interface. The pipeline improves signal-to-noise, contrast, and spatial resolution before performing accurate detection and segmentation of spine heads, spine necks, dendritic branches, and soma. Unlike existing tools, RESPAN operates through an intuitive graphical user interface (GUI), eliminating coding requirements while enabling analysis, model training in multiple environments, and validation. Additionally, it generates comprehensive readouts, including visual and tabulated data detailing dendritic spine morphology, signal intensity, spatial metrics as well as individual spine tracking for temporally longitudinal imaging. We demonstrate RESPAN’s high accuracy and robustness across multiple imaging modalities, representing a significant advancement in making advanced automated dendritic spine analysis accessible to the research community.

## RESULTS

### Development of RESPAN

The RESPAN pipeline (**Figure 1**) integrates GPU image processing and multiple deep-learning approaches to enable high-throughput fluorescent image segmentation, 3D reconstruction, and analysis of dendritic branches and dendritic spines. By integrating content-aware restoration and 3D convolutional neural network segmentation, RESPAN improves spine detection accuracy and sensitivity to morphological variations, particularly in challenging experimental conditions such as in vivo two-photon microscopy and rapid volumetric imaging of large tissue volumes. To ensure broad accessibility, RESPAN is provided both as Python code and as a standalone Windows application with an intuitive, unified GUI, allowing users to run batch analysis, train models, and perform analysis validation all within the same interface. Combining these features and multiple software environments within one interface substantially increases utility and reduces barriers for researchers without programming expertise.

**Figure 1.**
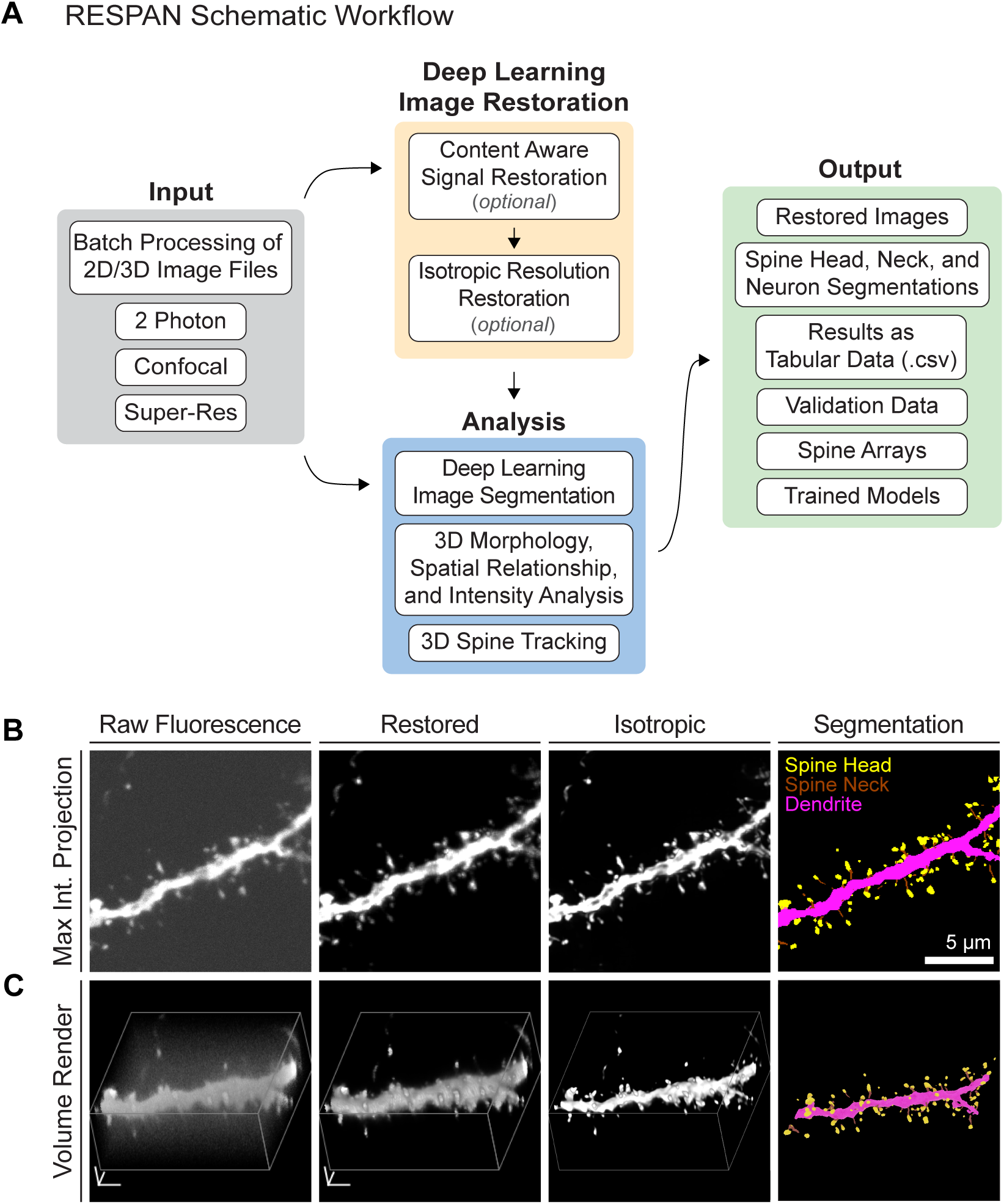
Overview of the RESPAN Workflow for Dendritic Spine Analysis (A) Schematic representation of the RESPAN workflow, beginning with the input of 2D/3D .tif image files and progressing through multiple stages including signal restoration, isotropic resolution restoration, deep learning image segmentation, and generation of quantitative outputs. RESPAN accommodates images from a variety of microscopy techniques including two-photon, confocal, and super-resolution microscopy. (B) Upper panels: Example of a raw fluorescence image acquired with a spinning disk confocal microscope, shown as a maximum intensity projection (MIP). (C) Lower panels: 3D renderings of the same dataset. Raw fluorescence images highlight reduced signal and resolution related to image acquisition. Following content aware image restoration, improvements in signal and contrast, essential for accurate spine detect, are apparent. Subsequent processing of the restored image to improve axial resolution reduces anisotropy, thus improving the accuracy of morphological measurements. Finally deep-learning-based segmentation on the restored image provides precise labels for dendrites, spine heads, and necks. Lower panel: 3D view of the final segmentation output. Scale bars: 2 µm.

#### Step 1. Deep Learning Image Restoration (optional)

Prior to segmentation, RESPAN provides the option to perform image restoration to enhance signal quality and contrast. This is a critical step for accurate spine detection in challenging imaging conditions where phototoxicity and temporal resolution requirements limit image quality. To achieve this, RESPAN employs a 3D content-aware CSBDeep model ^22^, trained using paired low and high signal-to-noise ratio (SNR) image volumes. We acquired these image sets in various samples to best reflect the diversity of imaging conditions and biological features, including soma, dendritic branches, and spines, thus maximizing the generalizability of the trained model. This approach enables high-quality restoration that preserves important biological features. For spinning disk confocal microscopy, we used a set of six paired image volumes with dimensions 1500 x 1500 x 130, with a pixel size of 65nm and a Z-step size of 150nm. Prior to training, the high SNR datasets were processed using a blind deconvolution. For 2- photon microscopy, we used 30 paired image volumes with average dimensions ∼300 x 600 x 12, with a pixel size of 102nm and a Z-step size of 1µm. For both models, training was performed in Python using image augmentation (rotation, mirroring, and flipping) with patch sizes of 64 x 64 x 16 for 100 epochs as previously described parameters ^22^. Representative examples of CARE restoration applied to low-SNR datasets can be seen in **Figure S1**, illustrating how this approach can improve fluorescence signal while preserving morphological details crucial for downstream spine segmentation.

Next, an optional axial restoration approach is used to reduce anisotropy and improve the accuracy of spine morphology measurements. This step is particularly important for conventional microscopy techniques like confocal microscopy, where the axial resolution can be 2-3 times worse than the lateral resolution, and even more critical for two-photon microscopy, where the disparity between lateral and axial resolution can be 4-5 times or greater due to the elongated point spread function. These resolution differences significantly impact the accuracy of 3D measurements. To address such challenges, RESPAN incorporates Self-Net. This two-stage unsupervised neural network approach leverages the natural anisotropy of microscopy data to improve axial resolution without requiring paired training data ^23^. By using lateral images as rational targets, Self-Net can learn to enhance axial resolution while maintaining high fidelity to genuine biological features. This approach necessitates models trained on image data that match the same specific experiment and imaging conditions for accurate results. To facilitate model training using user-specific datasets, we have integrated this capability within the RESPAN GUI using code adapted from the original implementation of Self-Net ^23^.

#### Step 2. Image segmentation

For image segmentation, RESPAN employs a 3D full-resolution nnU-Net model, a self-configuring framework for biomedical image segmentation ^24^. This approach automatically optimizes network architecture and training parameters based on input data properties, ensuring robust performance across varied imaging conditions. To ensure broad utility, we created a diverse training dataset comprising 47 image volumes containing 2,489 expert-annotated spines from multiple imaging modalities commonly used in the field: Zeiss Airyscan confocal microscopy (63x 1.4 NA oil PlanApochromat objective), Yokogawa W1 spinning disk confocal microscopy (100x 1.35 NA silicone PlanApochromat objective), and Bergamo two-photon microscopy (Olympus 25x 1.0 NA SCALEVIEW-A2 Plan objective).

In addition to modality-specific models, we pooled images from different modalities to create a generalizable high-resolution model. This involved standardizing the training data by resampling the data from the Airyscan confocal datasets to match the spinning disk confocal datasets with a voxel size of 65 x 65 x 150nm. We expanded datasets through augmentation to further enhance model performance, including random rotations, flips, shot noise addition, and Gaussian blurring (**Figure S2**). This augmentation strategy improved model performance when combined with the on-the-fly patch augmentations generated by nnU-Net during training. Models were trained for 1000 epochs with an initial learning rate of 0.01 following established parameters ^24^. To train our best-performing Global Model 1, we leveraged nnUNet v2.4 on a single H100 GPU instance from the Lambda GPU Cloud. We used the Lambda Stack environment, which includes PyTorch and other dependencies for nnU-Net already compiled, enabling straightforward deployment on x86_64 architectures. Notably, we observed a marked improvement in segmentation accuracy after increasing the training batch size to 19 and the patch size to [32, 256, 256]. We managed the Python environment and dependencies on Ubuntu 22.04 LTS, using Lambda Stack’s preconfigured packages to streamline setup and execution. All relevant installation commands and hardware configuration details are provided in our public repository to promote reproducibility on multiple hardware platforms.

Validation of our high-resolution global model against ground truth labels demonstrated excellent performance, achieving Dice scores of 0.927 for spines, 0.986 for dendrites, and 0.991 for soma. For cases where high-resolution resampling may be undesirable due to increased memory requirements, we recommend using a modality-specific model matching the acquisition resolution. To maximize accessibility, RESPAN automatically scales input data to match model requirements while providing analysis output in the original resolution. This ensures that new users and novel datasets are likely to yield interpretable results without requiring additional preprocessing before using RESPAN.

#### Step 3. RESPAN output and quantification of dendritic spine morphometry

Following segmentation, RESPAN performs a comprehensive analysis of dendritic spines, providing both numerical measurements (**Tables 1** and **2**) and multiple validation outputs (**Figure S3**). Measurements include the volume of each spine, its centroid location, mean, minimum, and maximum intensity across available channels, and the distance from both the dendrite shaft and geodesic distance from the soma. Additional derived measurements, such as spine neck length and spine head width, are computed as previously described ^25^. Users can also specify constraints (e.g., minimum/maximum spine volume or distance from dendrite) to filter out spurious detections.

Summary statistics include per-dendrite and per-image measurements of spine number, density, dendrite length, and average spine measurements. RESPAN also supports the export of dendritic tracings for a single neuron or dendrite per image in standard .swc format via Vaa3D’s APP2 plugin ^26^, as well as 2D/3D visualization arrays of all segmented spines for rapid validation. For transparency, intermediate data (e.g., labeled segmentations, skeletons, and distance maps) can be saved, allowing users to verify segmentation accuracy and further refine segmentation models.

Importantly, RESPAN also includes a built-in validation module capable of automatically comparing results to ground truth annotations. This is essential for auditing RESPAN’s performance and validating and reporting the accuracy of new restoration and segmentation models and parameters when publishing with RESPAN. The metrics provided by the validation module are described in **Table 3**.

**Table 1.** Individual Spine Measurements.

| Column | Title | Contents |
| --- | --- | --- |
| 1 | spine_id | Unique identifier for each detected spine. |
| 2 | x | X-coordinate of the spine in image space (pixels). |
| 3 | y | Y-coordinate of the spine in image space (pixels). |
| 4 | z | Z-coordinate of the spine in image space (pixels). |
| 5 | dendrite_id | Identifier for the dendrite associated with the spine. |
| 6 | geodesic_dist | Geodesic distance from the spine to its parent dendrite ($\mu\text{m}$). |
| 7 | spine_area | Cross-sectional area of the entire spine ($\mu\text{m}^2$). |
| 8 | spine_vol | Volume of the entire spine ($\mu\text{m}^3$). |
| 9 | spine_vol_m | Measured spine volume adjusted for voxel resolution ($\mu\text{m}^3$). |
| 10 | spine_sa_m | Spine surface area adjusted for voxel resolution ($\mu\text{m}^2$). |
| 11 | spine_length | Total morphological length of the spine from the dendrite base to the spine head tip ($\mu\text{m}$). |
| 12 | head_area | Cross-sectional area of the spine head ($\mu\text{m}^2$). |
| 13 | head_vol | Volume of the spine head ($\mu\text{m}^3$). |
| 14 | head_vol_m | Measured volume of the spine head adjusted for voxel resolution ($\mu\text{m}^3$). |
| 15 | head_sa_m | Surface area of the spine head adjusted for voxel resolution ($\mu\text{m}^2$). |
| 16 | head_length | Length of the spine head, measured from the neck boundary to the head tip ($\mu\text{m}$). |
| 17 | neck_area | Cross-sectional area of the spine neck ($\mu\text{m}^2$). |
| 18 | neck_vol | Volume of the spine neck ($\mu\text{m}^3$). |
| 19 | neck_vol_m | Measured volume of the spine neck adjusted for voxel resolution ($\mu\text{m}^3$). |
| 20 | neck_sa_m | Surface area of the spine neck adjusted for voxel resolution ($\mu\text{m}^2$). |
| 21 | neck_length | Length of the spine neck, measured from the dendrite boundary to the head boundary ($\mu\text{m}$). |
| 22 | spine_length_dm | Alternative spine length measurement derived from distance mapping ($\mu\text{m}$). |
| 23 | spine_C1_int_density | Fluorescence integrated density in Channel 1 within the entire spine. |
| 24 | spine_C1_max_int | Maximum fluorescence intensity in Channel 1 within the entire spine. |
| 25 | spine_C1_mean_int | Mean fluorescence intensity in Channel 1 within the entire spine. |
| 26 | dist_to_dendrite_dm | Geodesic distance from the spine to the dendrite ($\mu\text{m}$). |
| 27 | dist_to_soma | Geodesic distance from the spine to the soma ($\mu\text{m}$). |
| 28 | head_C1_mean_int | Mean fluorescence intensity in Channel 1 within the spine head. |
| 29 | head_C1_max_int | Maximum fluorescence intensity in Channel 1 within the spine head. |
| 30 | head_C1_int_density | Fluorescence integrated density in Channel 1 within the spine head. |
| 31 | neck_C1_mean_int | Mean fluorescence intensity in Channel 1 within the spine neck. |
| 32 | neck_C1_max_int | Maximum fluorescence intensity in Channel 1 within the spine neck. |
| 33 | neck_C1_int_density | Fluorescence integrated density in Channel 1 within the spine neck. |

**Table 2.** Detected Spines Summary.

| Column | Title | Contents |
| --- | --- | --- |
| 1 | Filename | Name of the analyzed image file. |
| 2 | res_XY | Lateral (XY) resolution of the image in micrometers per pixel ($\mu\text{m}/\text{pixel}$). |
| 3 | res_Z | Axial (Z) resolution of the image in micrometers per pixel ($\mu\text{m}/\text{pixel}$). |
| 4 | dendrite_id | Unique identifier for the segmented dendrite. |
| 4 | dendrite_length | Total length of the segmented dendrite, in micrometers ($\mu\text{m}$). |
| 5 | dendrite_vol | Total dendritic volume, in cubic micrometers ($\mu\text{m}^3$). |
| 6 | total_spines | Total number of detected spines along dendrite. |
| 7 | spines_per_um | Number of spines per micrometer of dendrite length (spines/$\mu\text{m}$). |
| 8 | spines_per_um3 | Number of spines per cubic micrometer of dendrite volume (spines/$\mu\text{m}^3$). |
| 9 | avg_spine_area | Mean cross-sectional area of the detected spines, in square micrometers ($\mu\text{m}^2$). |
| 10 | avg_spine_vol | Mean volume of the detected spines, in cubic micrometers ($\mu\text{m}^3$). |
| 11 | avg_spine_vol_m | Mean mesh volume of spines ($\mu\text{m}^3$). |
| 12 | avg_spine_sa_m | Mean surface area of spines ($\mu\text{m}^2$). |
| 13 | avg_spine_length | Mean length of the spines, in micrometers ($\mu\text{m}$). |
| 14 | avg_head_area | Mean cross-sectional area of the spine heads ($\mu\text{m}^2$). |
| 15 | avg_head_vol | Mean volume of the spine heads ($\mu\text{m}^3$). |
| 16 | avg_head_vol_m | Mean mesh volume of spine heads ($\mu\text{m}^3$). |
| 17 | avg_head_sa_m | Mean surface area of spine heads ($\mu\text{m}^2$). |
| 18 | avg_head_length | Mean length of the spine heads ($\mu\text{m}$). |
| 19 | avg_neck_area | Mean cross-sectional area of the spine necks ($\mu\text{m}^2$). |
| 20 | avg_neck_vol | Mean volume of the spine necks ($\mu\text{m}^3$). |
| 21 | avg_neck_vol_m | Mean mesh volume of spine necks ($\mu\text{m}^3$). |
| 22 | avg_neck_sa_m | Mean surface area of spine necks ($\mu\text{m}^2$). |
| 23 | avg_neck_length | Mean length of the spine necks, in $\mu\text{m}$. |
| 24 | avg_spine_length_dm | Mean spine length derived from distance-map or similar metric, in $\mu\text{m}$. |
| 25 | avg_spine_C1_int_density | Mean integrated density in Channel 1 within all spines. |
| 26 | avg_spine_C1_max_int | Mean maximum fluorescence intensity in Channel 1 within all spines. |
| 27 | avg_spine_C1_mean_int | Mean fluorescence intensity in Channel 1 within all spines. |
| 28 | avg_dist_to_dendrite_dm | Mean distance from spines to the dendrite shaft, in $\mu\text{m}$. |
| 29 | avg_dist_to_soma | Mean distance from spines to the soma, in $\mu\text{m}$. |
| 30 | avg_head_C1_mean_int | Mean fluorescence intensity in Channel 1 within spine heads. |
| 31 | avg_head_C1_max_int | Maximum fluorescence intensity in Channel 1 within spine heads. |
| 32 | avg_head_C1_int_density | Mean integrated density in Channel 1 within spine heads. |
| 33 | avg_neck_C1_mean_int | Mean fluorescence intensity in Channel 1 within spine necks. |
| 34 | avg_neck_C1_max_int | Maximum fluorescence intensity in Channel 1 within spine necks. |
| 35 | avg_neck_C1_int_density | Mean integrated density in Channel 1 within spine necks. |

**Table 3.** Spine Analysis Summary (Temporal Analysis)

| Column | Title | Contents |
| --- | --- | --- |
| 1 | timepoint | Timepoint of the recorded data. |
| 2 | res_XY | Lateral resolution ($\mu\text{m}/\text{pixel}$). |
| 3 | res_Z | Axial resolution ($\mu\text{m}/\text{pixel}$). |
| 4 | dendrite_length | Total length of the segmented dendrite ($\mu\text{m}$). |
| 5 | dendrite_vol | Total dendritic volume ($\mu\text{m}^3$). |
| 6 | total_spines | Total number of detected spines at this timepoint. |
| 7 | spines_per_um | Density of spines per unit dendritic length (spines/$\mu\text{m}$). |
| 8 | new_spines | Number of newly detected spines at this timepoint. |
| 9 | pruned_spines | Number of spines that disappeared compared to the previous timepoint. |
| 10 | spines_per_um3 | Density of spines per unit dendritic volume (spines/$\mu\text{m}^3$). |
| 11 | avg_x | Mean x-coordinate of detected spines (pixels). |
| 12 | avg_spine_area | Mean spine cross-sectional area (pixels<sup>2</sup>). |
| 13 | avg_spine_area_um2 | Mean spine cross-sectional area ($\mu\text{m}^2$). |
| 14 | avg_spine_vol | Mean spine volume ($\mu\text{m}^3$). |
| 15 | avg_C1_mean_int | Mean fluorescence intensity in Channel 1 within spines. |
| 16 | avg_spine_length | Mean spine length ($\mu\text{m}$). |
| 17 | avg_C1_max_int | Maximum fluorescence intensity in Channel 1 within spines. |
| 18 | avg_dist_to_dendrite | Mean geodesic distance of spines to the dendrite ($\mu\text{m}$). |
| 19 | avg_dist_to_soma | Mean geodesic distance of spines to the soma ($\mu\text{m}$). |
| 20 | avg_C1_int_density | Mean fluorescence intensity density in Channel 1 within spines. |

### Comprehensive validation of RESPAN performance

We employed a comprehensive validation strategy combining both pixel-level and object-level metrics to evaluate RESPAN’s performance, following established practices in deep learning-based microscopy analysis ^22,24^. For pixel-level assessment, we measured the Intersection over Union (IoU) (|A ∩ B|/|A ∪ B|) and the Dice Coefficient (DC) (2|A ∩ B|/|A| + |B|), where A represents RESPAN output pixels and B represents ground truth pixels. These metrics proved particularly valuable for assessing the precise delineation of spine and dendrite boundaries. For object-level validation, to assess for accurately counting and characterizing individual spines, we employed an IoU threshold of 50% to quantify successfully detected objects. The object-level comparison generated true positives (TP), false positives (FP), and false negatives (FN), which we used to calculate two key metrics: the proportion of correctly detected objects (Precision = TP/(TP+FP)) and the proportion of true objects that were correctly detected (Recall = TP/(TP+FN)). Spine detection demonstrated excellent performance, achieving Precision and Recall scores of 0.988 and 0.9, respectively. We achieved a final F1 score of 0.94.

### Benchmarking RESPAN accuracy against human expert annotations

To assess the accuracy and reliability of RESPAN in detecting dendritic spines, we conducted a comprehensive comparison with manual annotations from multiple users of varying expertise (**Figure 2**). Using spinning disk confocal microscopy, we first acquired five independent fluorescence images of dendritic segments (**Figure 2A**). Two experts with years of experience in imaging dendritic spines and manual spine analysis carefully annotated these images to create a ground truth (GT) consensus dataset (**Figure 2B-2C**). We then used these GT datasets to evaluate spine detection performance against (1) four independent users with varying levels of expertise and (2) RESPAN output of the same images.

**Figure 2.**
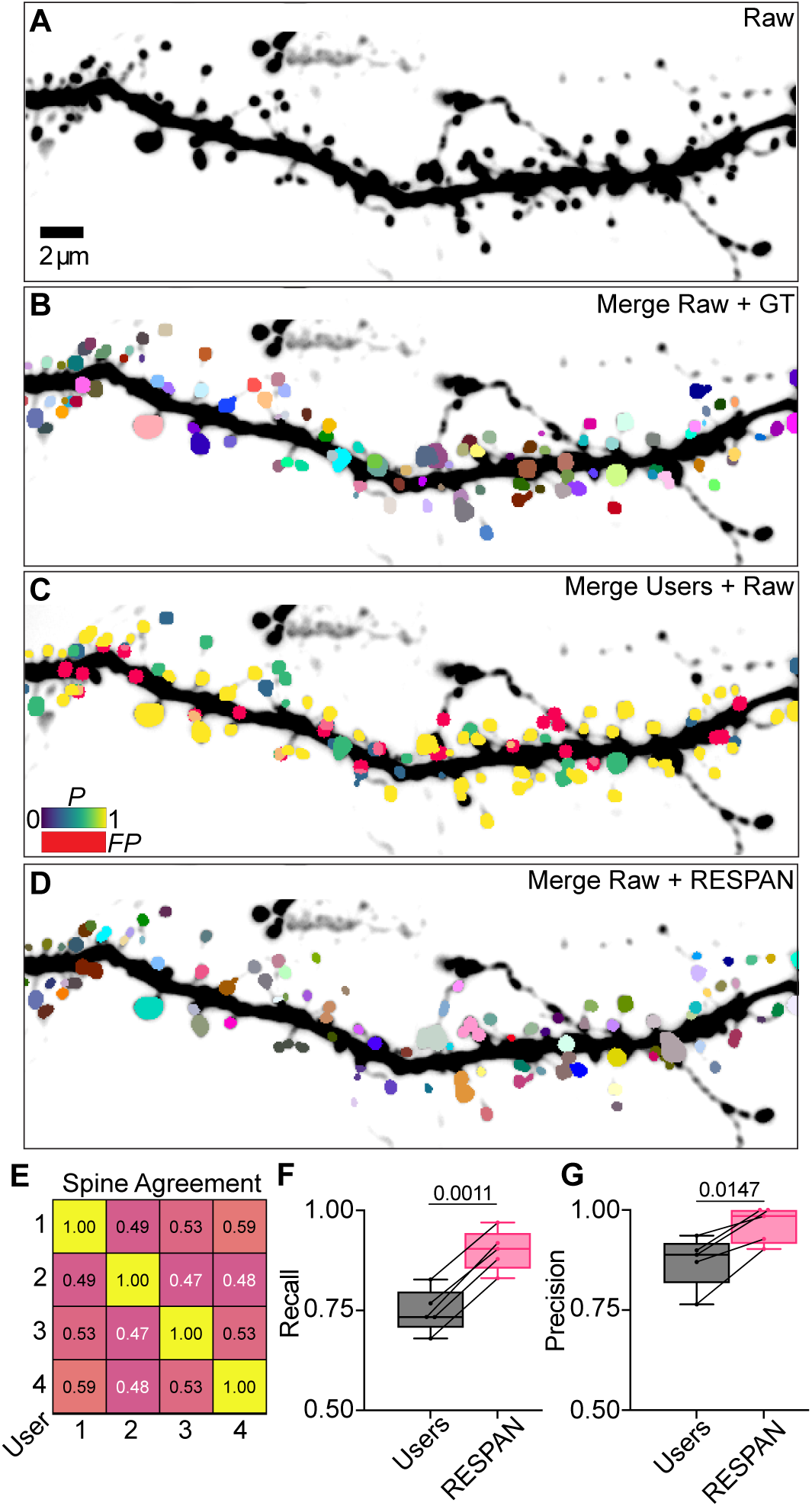
Evaluation of Multi-User Spine Detection Variability against RESPAN. (A) Raw fluorescence image of a dendritic segment used as input for both manual annotation by multiple experts and automated processing by RESPAN. (B) Ground truth (GT) annotations determined by a consensus of experienced researchers; colors identify unique spine. (C) Composite image merging the raw fluorescence with the GT annotations overlaid. Spine colors reflect the probability of detection per spine from four independent users, with false positive detections labeled in red. (D) Composite image merging raw fluorescence with RESPAN spine detections, with colors reflecting every unique spine identified via a connected component analysis. (E) Inter-user correlation matrix showing the pairwise detection agreement among human annotators, with values ranging from 0 (no agreement) to 1 (perfect agreement). (F-G) Box plots comparing recall and precision between human annotators and RESPAN, showing that RESPAN consistently achieves higher recall and precision, reflecting improved sensitivity and fewer false positives relative to manual human annotation. Pairwise detection agreement among human annotators (values from 0 to 1) was quantified using Pearson correlation. Recall and precision were compared between human annotators and RESPAN using a paired two-tailed t-test (n = 5 pairs) at the 90% confidence interval. The resulting p-values were p = 0.0011 for recall (F) and p = 0.0147 for precision (G). p < 0.05 was considered significant.

Each user analyzed the 3D raw fluorescence images in Fiji, marking putative spines at their perceived center of mass using the multi-point tool. We compiled the ROIs placed by each user and superimposed them onto the GT consensus mask. We used custom Python code to quantify the number of multi-point ROIs detected for each GT spine across all users and images, generating a comprehensive spine probability map (**Figure 2C**). This representation illustrates visually the significant variability in users’ ability to identify spine heads in a concordant manner.

To determine inter-user agreement, we generated a correlation matrix depicting pairwise detection agreement among users, with values ranging from 0 (no agreement) to 1 (perfect agreement) (**Figure 2E**). The analysis revealed that inter-user agreement on spine detection remained consistently low, suggesting potential statistical biases in manual annotation approaches. Further analysis examined the probability distribution of spines being detected by varying numbers of users: no users (0), one user (0.25), two users (0.50), three users (0.75), or all four users (1.00) (**Figure 2C**).

We then processed these same fluorescence images through the RESPAN pipeline for automated spine segmentation. Using custom Python code, we then overlaid RESPAN’s segmentation output onto our GT consensus mask, employing an IoU criterion of 0.50 to determine correct spine detection. Comparing RESPAN’s performance to user detection probabilities revealed two key findings: First, RESPAN showed a marked decrease in missed detections compared to human annotators, more accurately identifying spines present in the GT consensus mask but missed by multiple users. Second, RESPAN demonstrated improved detection of spines identified by only a subset of users, indicating superior sensitivity in challenging cases.

These results demonstrate that RESPAN outperforms manual spine identification while alleviating inter- individual variability and human observer bias in image annotation. Furthermore, the automated approach provides comprehensive morphological readouts that are not feasible with human annotation, enabling a more thorough and unbiased analysis of dendritic spine populations.

### Demonstrating the importance of image restoration in spine detection

We next investigated how content-aware restoration could improve RESPAN’s segmentation performance in low-SNR imaging scenarios commonly encountered in live or in vivo two-photon experiments (**Figure 3**). Although our Global Model 1 already provides reasonable segmentation on noisy datasets, our results indicate that applying a CARE restoration model (Weigert et al., 2018) prior to RESPAN segmentation further enhances spine detection under these challenging conditions. Under these optimized training conditions, CARE substantially improved spine detection and segmentation metrics when compared to low-SNR data (**Figure 3D–G**). Notably, in this study, we deliberately used extremely low-SNR images to reflect a “worst-case” scenario. Even under these stringent conditions, the detection of spines with IoU ≥ 0.5 rose from a median of 0.70 to 0.75, and the fraction of true-positive spines increased from 0.76 to 0.82. Furthermore, the F1 score, which captures both recall and precision, showed a marked increase in its cumulative distribution as the IoU threshold became more stringent, reflecting the improved robustness of restored segmentations.

**Figure 3.**
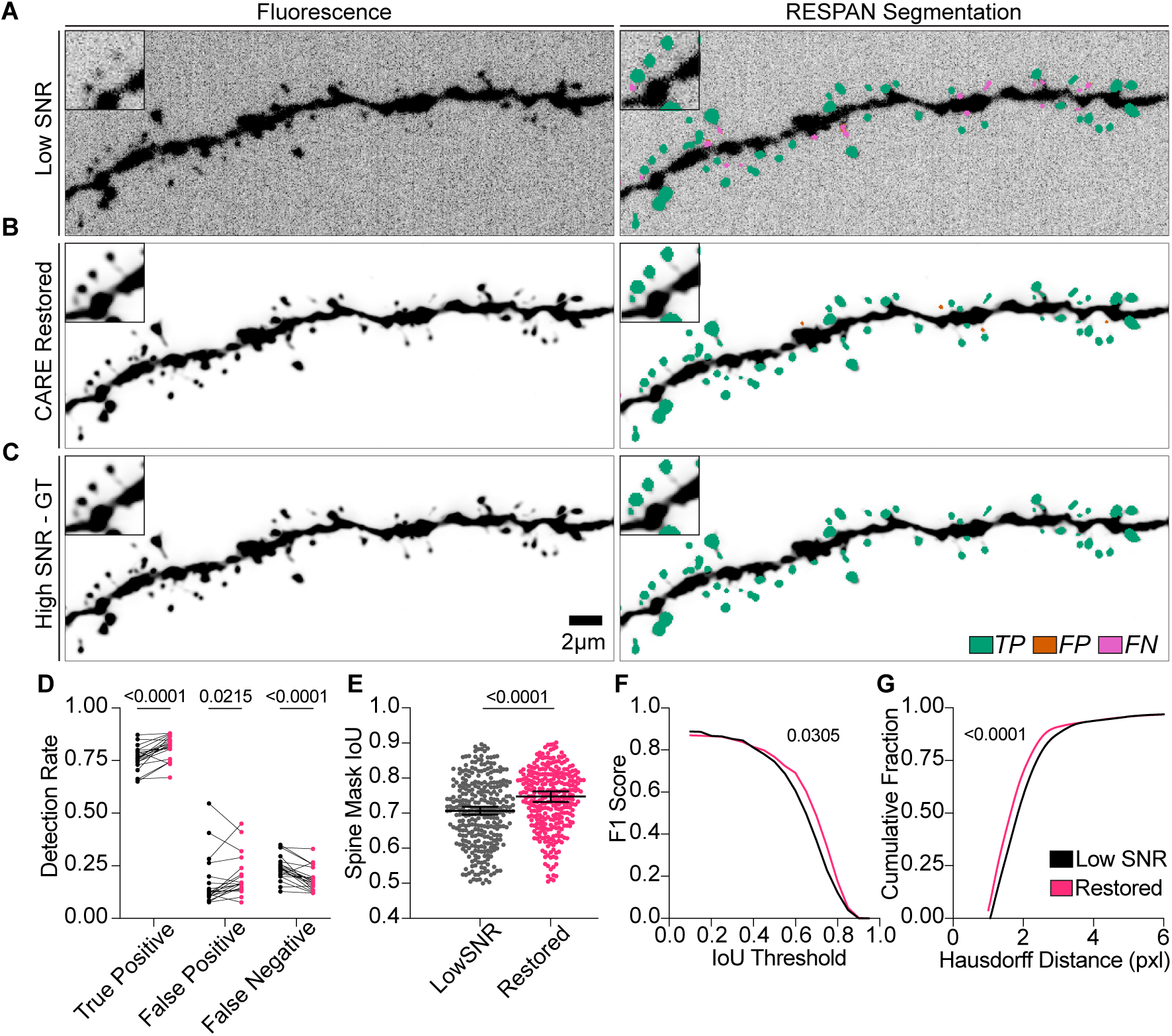
Importance of Image Restoration in Spine Detection Accuracy Using RESPAN. (A–C) Maximum-intensity-projection images of a dendritic segment acquired under low SNR conditions (A), following CARE restoration (B), and at high SNR (C). The right panels in each row show RESPAN’s segmentation outputs, with color-coded spines denoting true positives (TP, green), false positives (FP, orange), or false negatives (FN, magenta). Insets highlight spines that are barely distinguishable in the low-SNR dataset but become visible after restoration. Scale bar, 2 µm. (D) Detection rates (TP, FP, and FN) for low-SNR versus restored datasets. Statistical comparisons were performed via Wilcoxon matched-pairs signed-rank tests with a two-stage step-up method (Benjamini, Krieger, Yekutieli) to control the false discovery rate. (E) Spine mask intersection-over-union (IoU=0.5) for low-SNR and restored images. Each point represents a matched spine, and the line/whiskers depict the median and 95% CI. *p* values were obtained using Wilcoxon matched-pairs signed-rank tests. (F) F1 scores plotted across increasing IoU thresholds (0.1–0.9). Comparisons between low-SNR (black) and restored (magenta) curves were performed using Wilcoxon matched-pairs signed-rank tests. (G) Cumulative distributions of Hausdorff distances for low-SNR (black) and restored (pink) spines, where lower values indicate closer alignment with the ground-truth spine shape. Data were analyzed by a Kolmogorov–Smirnov test, revealing significantly reduced Hausdorff distances in restored data. All *p* values are displayed as integers within the figure.

We additionally assessed the boundary accuracy using the Hausdorff distance (**Figure 3G**), which measures how far the most extreme mismatched boundary point lies from ground-truth surfaces. After CARE restoration, we observed significantly lower Hausdorff distances, indicating that even at the “worst- matching” boundary points, predicted spines were closer to their ground-truth counterparts. This improvement is crucial for downstream measurements of spine volume or surface area, where a single poorly placed boundary point could skew morphological calculations.

Equally important, the ability to reliably restore low-SNR data has practical benefits for experimental design. By capturing images at lower exposure times and reduced laser power, users can shorten their time at the microscope while mitigating photobleaching and phototoxicity. In turn, the restored images effectively regain high-SNR features, allowing users to move forward with accurate quantification. Thus, our workflow enables high-fidelity segmentation of spines with minimal imaging burden, which is particularly valuable for long time-lapse series or large-volume acquisitions, where preserving sample health and reducing acquisition time are paramount.

### Biological validation of RESPAN using established phenotypes affecting spine density and morphology

To validate RESPAN’s performance, we performed a blinded genotype/phenotype analysis using *in vivo* 2-photon microscopy of dendritic segments from layer 2/3 pyramidal neurons (PNs) of WT and SRGAP2^+/-^knockout mice. We wanted to determine if RESPAN would detect differences in spine density and morphology previously well-established. Our lab demonstrated that adult layer 2/3 PNs from SRGAP2^+/-^ mice exhibit unique spine phenotypes: increased spine density, increased spine neck length but no significant change in spine volume distribution compared to adult WT littermates ^1,27–30^. For this analysis, neurons were sparsely labeled via *in utero* electroporation (IUE) with a plasmid FLEX-mGreenLantern together with low amount of another plasmid expressing Cre-recombinase ^31^. Imaging of individual dendritic segments of layer 2/3 PNs was performed *using in* vivo 2-photon microscope through a cortical window to capture spine morphology in live tissue (**Figure 4A**).

**Figure 4.**
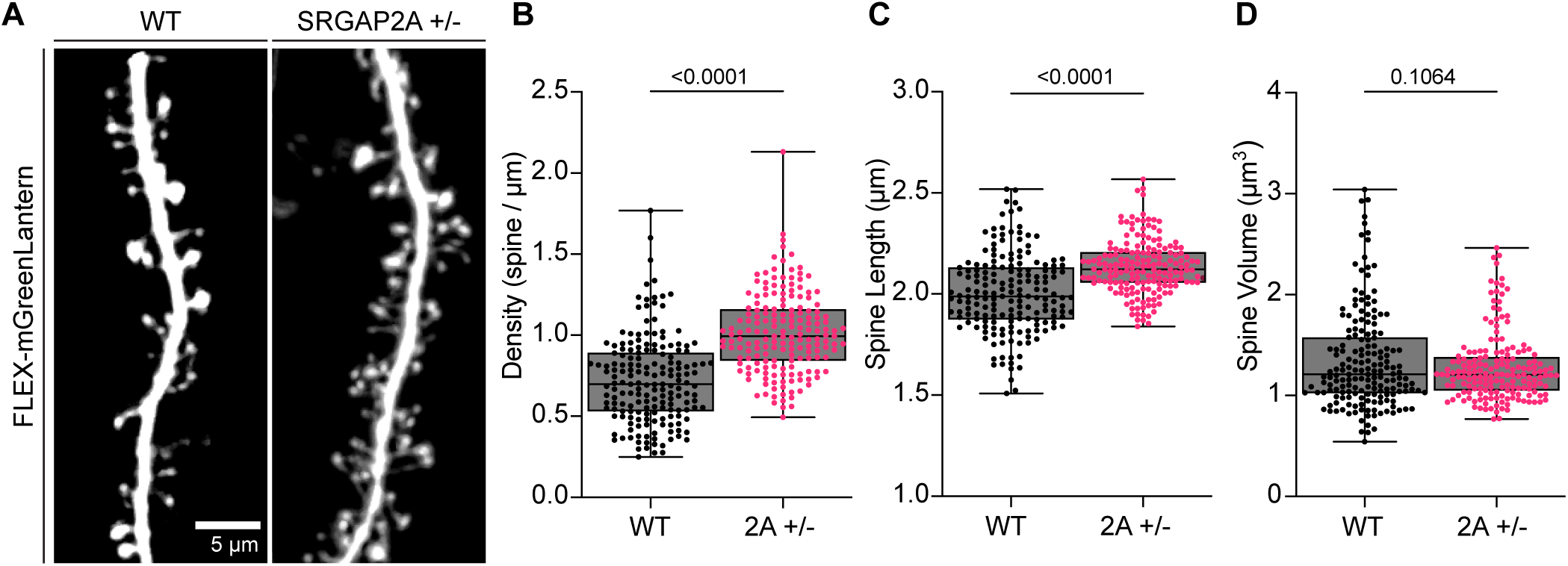
Blind Genotype/Phenotype Validation Using RESPAN. (A) *In vivo* 2-photon microscopy images of dendritic segments from wild-type (WT) and SRGAP2A +/- mutant neurons. Neurons were sparsely labeled through *in utero* electroporation (IUE) using low amounts of Cre recombinase and a FLEX-mGreenLantern construct, imaged through a cortical window to capture spine morphology. Scale bar, 2 µm. (B) Spine density comparison between WT and SRGAP2A +/- mutants, showing a significant increase in spine density per unit length in mutants. Results analyzed using the non-parametric Kolmogorov-Smirnov test (****p < 0.0001). (C) Spine length comparison between WT and SRGAP2A +/- mutants, with mutants exhibiting significantly longer spines. Analysis performed using the non-parametric Kolmogorov-Smirnov test (****p < 0.0001). (D) Comparison of spine volume between WT and SRGAP2A +/- mutants, revealing no significant difference in spine volume, as determined by the non-parametric Kolmogorov-Smirnov test (ns, not significant).

RESPAN analysis successfully replicated these established phenotypes through an unbiased and blinded approach. Quantitative analysis revealed that SRGAP2^+/-^ mutants displayed significantly higher spine density than WT neurons (**Figure 4B**, ****p < 0.0001), confirming the expected increase ^1,27–30^. Similarly, dendritic spines had significantly longer necks in SRGAP2^+/-^ mutants compared to WT neurons (**Figure 4C**, ****p<0.0001), matching our previous findings using manual analysis. Notably, there was no significant difference in spine volume between WT and SRGAP2^+/-^ mice (**Figure 4D**, ns), consistent with our previous results using manual measurements. This validation demonstrates RESPAN’s ability to accurately detect and quantify dendritic spine phenotypes in an unbiased manner, providing results that directly align with previously published evidence.

To further validate RESPAN’s quantification accuracy against established metrics, we analyzed spine morphology distributions in a larger population of WT neurons. We compared our results to previously published ground truth data where all excitatory and inhibitory spines on a single neuron were mapped and analyzed, providing a ground truth for spine distribution (**Figure 5A** and **5B**; ^1^). This analysis of 81,604 spines revealed characteristic distributions: spine length following a normal distribution and spine volume showing a Poisson distribution with predominant small spines and a smaller population of large spines. Using RESPAN, we analyzed 10,945 spines from WT neurons imaged by spinning disk confocal microscopy, applying spine volume thresholds of 0.035 µm³ and 2 µm³ (**Figure 5C** and **5D**). The results demonstrated that RESPAN consistently replicates established biological distribution patterns for spine length and volume.

**Figure 5.**
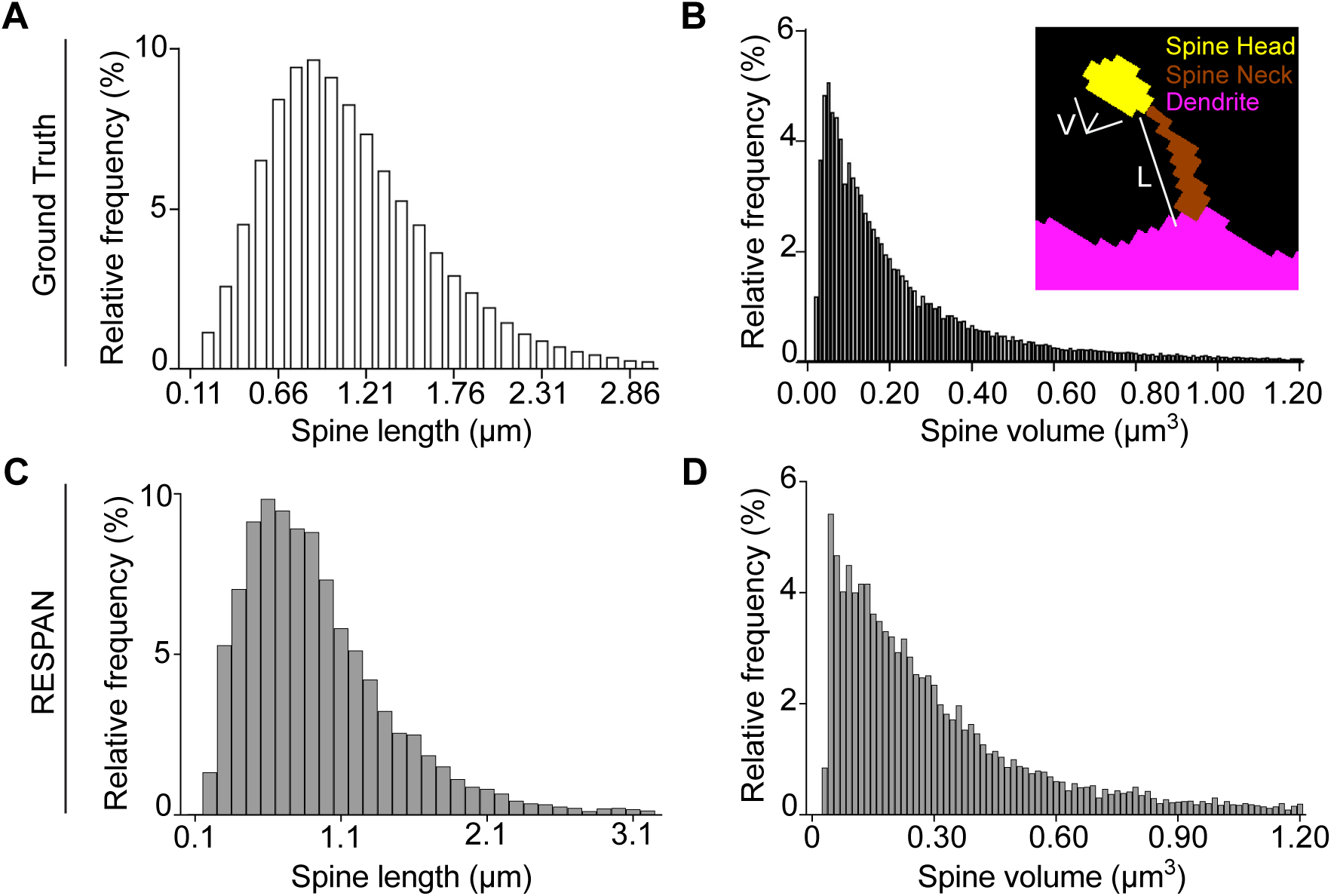
Measured Spine Lengths and Volumes Match Known Distributions Validation. (A) Histogram displaying the relative frequency distribution of spine lengths in wild-type (WT) neurons, based on a previous analysis of 81,604 spines, serving as ground truth. The x-axis represents spine length (µm) and the y-axis shows the percentage of spines within each length range. (B) Histogram showing the relative frequency distribution of spine volumes in WT neurons from the ground truth dataset. The x-axis represents spine volume (µm³) and the y-axis shows the percentage of spines within each volume range. The inset provides an example of a dendritic spine segment, color-coded to illustrate the spine head (yellow), spine neck (brown), and dendrite (magenta). Scale bar, 250 nm. (C) Histogram of spine length distributions in WT neurons based on RESPAN analysis of 10,945 spines. Results demonstrate a close match to ground truth distributions, highlighting RESPAN’s accuracy. (D) Histogram of spine volume distributions in WT neurons, showing a high degree of similarity to ground truth distributions. RESPAN analysis was performed on spinning disk confocal microscopy datasets, using spine head volume thresholds of 0.035 µm³ (minimum) and 2.0 µm³ (maximum). Output tables were merged to include all spines within these thresholds, with no post-processing modifications to the data.

### Spine tracking

Maintaining a consistent mapping of dendritic spines across multiple time points is a key challenge in longitudinal spine analysis, particularly under *in vivo* conditions where brain tissue may shift or deform. To address this, RESPAN provides a comprehensive spine tracking feature that integrates 3D volumetric registration, segmentation, and label matching into a single workflow. As depicted in **Figure 6**, we applied this method to two separate high-resolution two-photon Z-stacks obtained via a cranial window in a living mouse. RESPAN begins by allowing the user to perform either rigid or non-rigid 3D motion correction on each stack, ensuring that corresponding dendritic structures align accurately across sessions.

**Figure 6.**
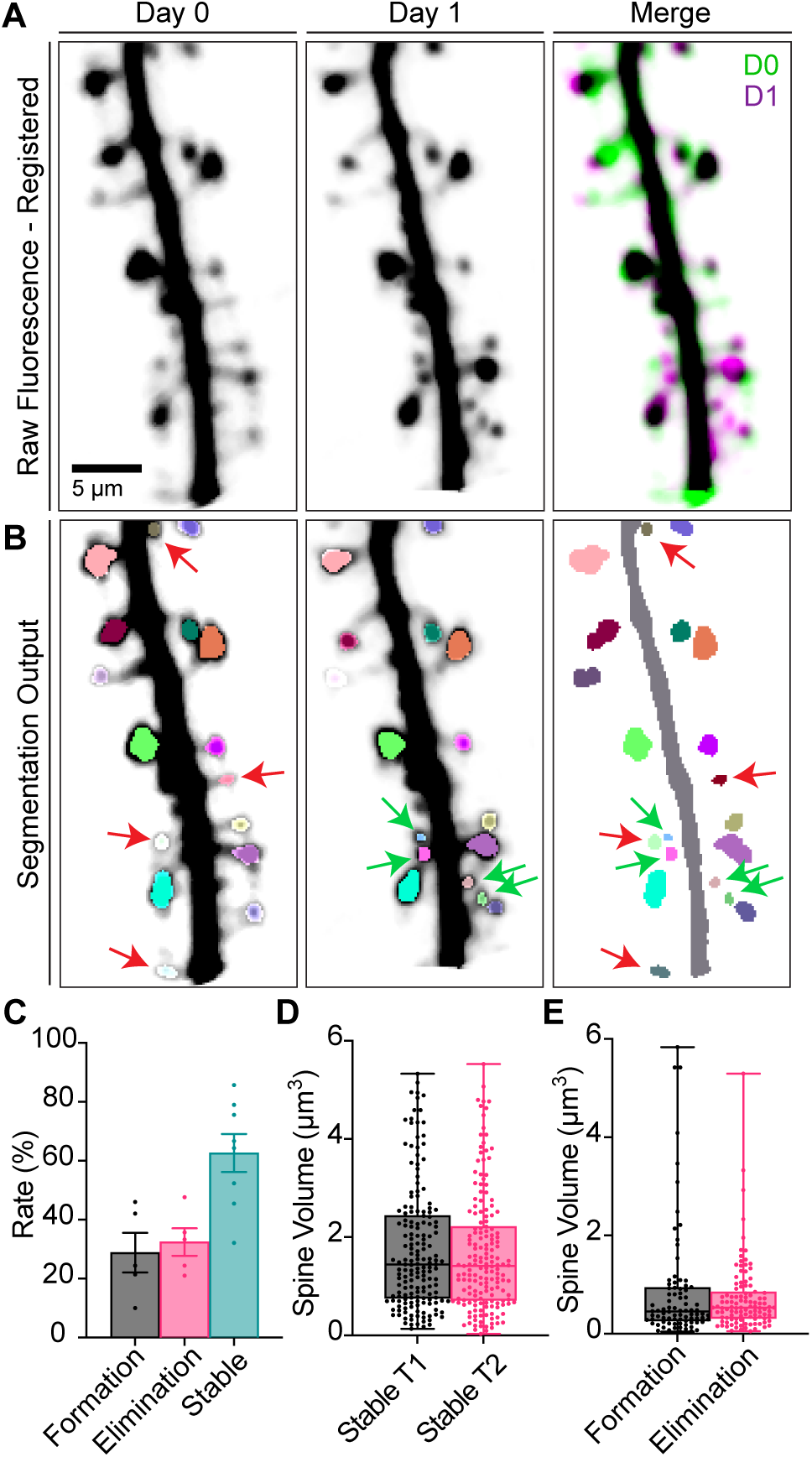
Application of RESPAN for the Analysis and Tracking of Spines *In Vivo*. (A) Representative two-photon microscopy images showing dendritic spines at Day 0 (left) and Day 1 (center), with their merged visualization (right). Raw fluorescence data was registered between timepoints to enable precise spine tracking. (B) Segmentation output from RESPAN, demonstrating accurate identification osf individual spines across time. Different colors indicate distinct spine categories: formation (green), elimination (red), and stable spines (same color between both timepoints). (C) Quantification of spine dynamics showing the rates (%) of stable spines across two timepoints (T1 and T2) and the relative proportions of spine formation and elimination events. Data are presented as mean ± SEM. (D) Distribution of spine volumes for newly formed and eliminated spines. (E) Volume comparison between stable spines at both timepoints. Box plots show median, quartiles, and distribution of individual measurements.

After registration, the pipeline proceeds with the segmentation of dendrites and spines. RESPAN then tracks these segmented objects across time, maintaining unique spine IDs: spines persisting between sessions share the same ID (and color), whereas newly formed and eliminated spines are labeled distinctly (**Figure 6A-B**). This color-coded visualization enables rapid identification of formation, elimination, and stabilization events (**Figure 6C-E**). Metrics evaluated during temporal analysis are documented in **Table 4**.

**Table 4.** Segmentation Validation Output.

| Column | Title | Contents |
| --- | --- | --- |
| 1 | Filename | Input image filename. |
| 2 | res_XY | Lateral image resolution ($\mu\text{m}/\text{pixel}$). |
| 3 | res_Z | Axial image resolution ($\mu\text{m}/\text{pixel}$). |
| 4 | gt_total_spines | Total spines in the ground truth image. |
| 5 | gt_total_spines_filtered | Total spines analyzed post-filtering in the ground truth image. |
| 6 | output_total_spines | Total spines detected by RESPAN. |
| 7 | output_total_spines_filtered | Total spines analyzed post-filtering by RESPAN. |
| 8 | TruePos IoU50 | True positive spines detected. |
| 9 | FalsePos IoU50 | False positive spines detected. |
| 10 | FalseNeg IoU50 | False negative spines. |
| 11 | Spine_precision IoU50 | Precision of spine detection. |
| 12 | Spine_recall IoU50 | Recall of spine detection. |
| 13 | gt_total_spine_vol | Total spine volume in the ground truth image ($\mu\text{m}^3$). |
| 14 | output_total_spine_vol | Total spine volume detected by RESPAN ($\mu\text{m}^3$). |
| 15 | total_spine_iou | IoU of spine voxels between ground truth and RESPAN. |
| 16 | total_spine_vol_difference | Difference in total spine volume between ground truth and RESPAN ($\mu\text{m}$). |
| 17 | gt_dendrite_length | Total dendritic length in the ground truth image ($\mu\text{m}$). |
| 18 | output_dendrite_length | Total dendritic length detected by RESPAN ($\mu\text{m}$). |
| 19 | dendrite_length_difference | Difference in dendritic length between ground truth and RESPAN ($\mu\text{m}$). |
| 20 | gt_dendrite_vol | Total dendritic volume in the ground truth image ($\mu\text{m}^3$). |
| 21 | output_dendrite_vol | Total dendritic volume detected by RESPAN ($\mu\text{m}^3$). |
| 22 | total_dendrite_iou | IoU of dendrite voxels between ground truth and RESPAN. |
| 23 | dendrite_vol_difference | Difference in dendritic volume between ground truth and RESPAN ($\mu\text{m}^3$). |

A unique strength of RESPAN is that it bundles this volumetric alignment and tracking capability in a user-friendly GUI, eliminating the need for specialized coding or computational expertise. This allows researchers to process folders containing multiple imaging sessions with their desired motion-correction method and readily verify alignment quality. To our knowledge, this marks the first GUI-based pipeline that combines 3D motion correction, restoration, segmentation, and longitudinal spine tracking in an integrated, semi-automated manner. By simplifying these previously complex workflows, RESPAN expands the practical feasibility of analyzing *in vivo* spine dynamics, enabling quantitative assessment of changes in spine number and morphology over time.

### Advantages and comparison with other methods

RESPAN addresses the critical challenges of reproducibility and efficiency in spine analysis, outperforming existing tools in both ease of use and accuracy (**Figure 7**). To benchmark its performance, we directly compared RESPAN’s spine detection results with those of DeepD3, a deep learning-based spine detection tool, and Imaris, a widely used commercial solution for 3D image analysis. Both, DeepD3 and Imaris exhibit higher rates of false positives (orange) and false negatives (magenta) than RESPAN (**Figure 7 A-D**).

**Figure 7.**
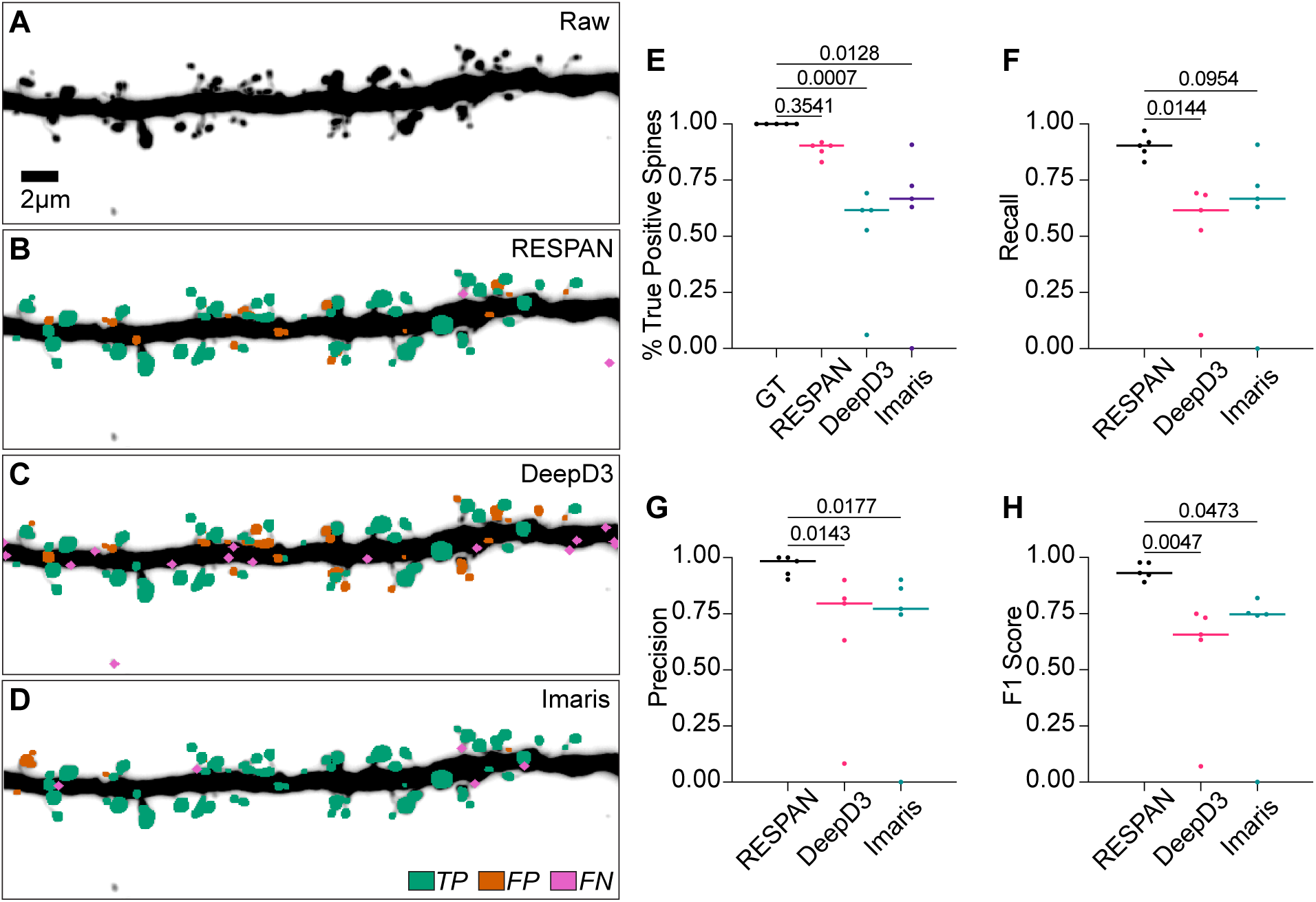
Comparison of RESPAN Performance with Other Software. (A) MIP of a raw fluorescence volume showing a dendritic segment. (B–D) Spine detection outputs from RESPAN (B), DeepD3 (C), and Imaris (D) overlaying the dendritic segment. Detected spines are color-coded as true positives (TP, green), false positives (FP, magenta), and false negatives (FN, orange). (E) Percentage of true positive spines detected by each method. (F) Recall scores for spine detection, with RESPAN demonstrating consistently higher recall, reflecting fewer false negatives. (G) Precision scores for spine detection, with RESPAN outperforming DeepD3 and Imaris by reducing false positive detections. (H) F1 scores for spine detection, representing the harmonic mean of precision and recall. RESPAN achieves superior F1 scores compared to other methods.

Quantitative metrics reinforce these observations: RESPAN achieves superior recall (p = 0.0144 vs. DeepD3; p = 0.0954 vs. Imaris), precision (p = 0.0143 vs. DeepD3; p = 0.0177 vs. Imaris), and F1 score (p = 0.0047 vs. DeepD3; p = 0.0473 vs. Imaris) (**Figure 7E–H**). The percentage of true positive (TP) spines detected by RESPAN also remains consistently higher than that of the other tools, reflecting more robust performance across different dendritic morphologies. These findings highlight RESPAN’s strengths as a fully automated, broadly applicable approach that is less sensitive to image-to-image variability and minimizes the need for manual calibration or correction.

For a broader overview of how RESPAN compares to other methods—including cost, supported platforms, and extensibility—please refer to **Table 5**. Whereas many commonly used workflows combine multiple software programs or rely on significant manual intervention, RESPAN integrates state-of-the- art deep learning modules for restoration, segmentation, and validation into a single GUI. This integration substantially reduces technical barriers and promotes reproducibility, allowing researchers to audit results and retrain models for new datasets seamlessly.

**Table 5.** Comparison of Existing Software Tools for Spine and Dendrite Analysis.

| Tool | RESPAN | SpineTool | DeepD3 | 3DSpan | DeepSpine Tool | Imaris |
| --- | --- | --- | --- | --- | --- | --- |
| Compatible Input Formats | .tif | .tif | .tif | .tif, .hdr, .img | .tif | .ims |
| Machine Learning Capabilities | Image restoration (SNR, resolution), Segmentation (2D/3D U-Net) | – | Segmentation (2D U-Net) | – | Segmentation (2D U-Net) | Segmentation (pixel classification) |
| Image Outputs | Spine, dendrite, soma masks (.tif), 2D/3D spine arrays (.tif), dendrite tracing (.swc) | Binarized spine image (.tif), spine meshes (.off) | Spine ROIs (.rois), spine masks (.tiff), prediction maps (.prediction) | Segmented sub-volumes (.hdr/.img) | Segmentation mask (.tif) | Objects visible within Imaris and can be converted to image data for export |
| Tabular Outputs | Spine counts and metrics (.csv) | Spine/dendrite classification (.json) | None | Segmentation analysis (.csv) | None | Selected parameters can be manually exported |
| Installation Requirements | Windows, NVIDIA GPU, Python 3.9, Anaconda | Windows, 1GB RAM, Anaconda | Python 3.7, Anaconda, TensorFlow | Windows | Windows, Python 3.6, CUDA 10.1, 8GB GPU | Windows/Mac |
| Training Data Acquisition Modality | 2-photon, spinning disk confocal, Airyscan | Confocal | 2-photon | 2-photon | – | – |
| User Interface | GUI (single/batch processing) | Jupyter Notebook | GUI (single image, scripting for batch) | GUI | GUI | GUI |
| Open Source | Yes | Yes | Yes | Yes | No | No |
| Batch Processing | Yes | No | Yes | No | No | Yes |
| User-Trainable Models | Yes | No | Yes | No | No | No |
| Spine Quantification | Yes | No | Yes | Yes | No | Yes |
| Validation Capability | Yes | No | Yes | No | No | No |
| ImageJ Compatibility | Yes | Yes | Yes | No | Yes | No |

To our knowledge, RESPAN is the only freely available tool that integrates multiple state-of-the-art deep learning approaches - content-aware image restoration ^22^, axial resolution enhancement ^23^, and self-configuring image segmentation ^24^. While recent advances have shown the power of individual deep learning approaches for specific tasks, such as DeepD3 for spine detection or CARE for image restoration, combining these capabilities requires significant computational expertise and manual data handling between different software tools. RESPAN addresses this limitation by providing a complete, integrated solution that maintains high performance while eliminating the technical barriers and overhead traditionally associated with using multiple separate analysis tools. This integration is valuable for ensuring reproducibility and maintaining data provenance throughout the analysis pipeline. Additionally, instead of performing image operations on the CPU, RESPAN leverages GPU processing to efficiently handle images in parallel, including in the rapid generation of 2D and 3D spine arrays.

Critically, unlike other methods, RESPAN provides a built-in tool for automatically validating the quality of analysis against ground-truth data. This addresses an unmet need, as segmentation validation is often performed manually or less rigorously by researchers with limited coding experience. By incorporating this capability, we believe RESPAN can encourage a rigorous approach to automated quantification and increase the reproducibility of published results. RESPAN also facilitates training new restoration and segmentation models directly within the GUI, which would otherwise require separate Python environments and coding experience, significantly reducing barriers to adopting this approach.

RESPAN significantly lowers the barriers to performing advanced image quantification for researchers without coding expertise. This addresses challenges presented by other tools that require a combination of Python or MATLAB scripts without a GUI, including tools that have been incorporated into our approach. Furthermore, unlike other methods that limit analysis outputs to dendritic spine coordinates ^9^ or require the use of multiple programs not within the same environment ^9^, our approach provides comprehensive 3D analysis within a single graphical user interface environment. RESPAN can also be readily adapted to new research questions by correcting initial results from the provided models using Fiji ^32^, a widely used image analysis software, and the tools within RESPAN.

## DISCUSSION

Dendritic spine analysis remains a cornerstone for understanding neuronal connectivity. Yet, current approaches—predominantly manual or semi-automated—are constrained by interactive annotation steps, image-specific parameters, user-dependent variability, and limited throughput. In addressing these challenges, RESPAN introduces a fully integrated, deep learning-enhanced solution that significantly broadens the scope and reliability of automated spine analysis.

One of RESPAN’s significant contributions lies in its integrated architecture, consolidating image restoration, axial resolution enhancement, and 3D segmentation into a single application. This design minimizes biases intrinsic to manual methods and facilitates reproducible and direct comparisons across diverse imaging modalities. With increasing focus on methodological rigor, RESPAN provides a streamlined and robust framework for quantifying synaptic plasticity, neuronal development, and disease models.

RESPAN also addresses significant barriers related to accessibility. Harnessing state-of-the-art computational image analysis has historically required specialized expertise or costly commercial platforms. By integrating GPU acceleration, model training, and analysis validation into a user-friendly interface, RESPAN lowers these barriers and democratizes the use of advanced analysis tools. This broad accessibility has the potential to accelerate discoveries in fields ranging from basic synaptic physiology to translational research on neurodevelopmental disorders and neurodegenerative diseases, areas where subtle changes in spine morphology and distribution may reveal profound shifts in neuronal function.

Looking forward, RESPAN’s architecture provides a platform for future extensions. Further integration of segmentation models and analysis tailored to related biological features, such as mitochondria and organelle analysis along dendrites and in relation to spines, as well as pre- and post-synaptic protein analysis, will further enhance its utility.

In conclusion, RESPAN represents a significant advancement in automated dendritic spine analysis, delivering accurate, robust, and accessible quantification to the neuroscience community. By enabling unbiased, high-throughput measurements of spine morphology and distribution, RESPAN paves the way for more comprehensive and reproducible studies of neuronal connectivity. By sharing the restoration and segmentation models along with the code as an open-source platform, RESPAN fosters collaboration and iterative development, allowing it to evolve alongside the research community’s needs and ensuring continued relevance and impact.

## Limitations of the study

Despite these advantages, RESPAN does have limitations. The pipeline’s performance depends heavily on training datasets that faithfully represent the biological and imaging conditions of interest. While RESPAN includes tools for model retraining, future iterations could benefit from more comprehensive pre-trained models covering a wider range of biological and imaging conditions.

RESPAN currently only supports input data in TIFF format, requiring conversion of proprietary formats to TIFF before use in our software. To overcome this limitation, we provide a Fiji macro to facilitate batch conversation to TIFF using OMERO Bio-Formats ^33^. We also generate our output data in TIFF for ease of use, but future releases would benefit from added support for the Zarr format to improve scalability.

Users should be aware of the challenges of applying deep learning models to microscopy data ^22,34^. Content-aware restoration and axial enhancement models should ideally be trained on data matching the specific biological features, resolution, and imaging modality of the intended application. When models are applied to data significantly different from their training set, they may perform sub-optimally and introduce artifacts by hallucinating features or degradation of genuine structures. This is particularly relevant when comparing experimental conditions with substantial morphological differences - in such cases, training data should encompass the full range of expected biological variation to ensure reliable results. Furthermore, as demonstrated in recent studies ^23^, the performance of these models can degrade when the resolution anisotropy ratio exceeds 4-fold or when input data has extremely low signal-to-noise ratios. To address these limitations, RESPAN includes model validation and retraining tools, allowing users to optimize performance for their specific experimental conditions.

## RESOURCE AVAILABILITY

### Lead contact

Luke A. Hammond.

### Data and code availability

● RESPAN code and executable available for download at: https://github.com/lahammond/RESPAN

`o` RESPAN is provided as Python code and as a standalone Windows application with a graphical user interface (GUI), eliminating the need for programming expertise.
● Five example spinning disk confocal datasets and expected results generated by RESPAN are available for download at:

`o` https://drive.google.com/drive/folders/1E8vJ4vUvrhaIZ0h7o1QyjzLW3YaTMaTJ?usp=sh <u>aring</u> (Google link provided in lieu of Mendeley repository currently awaiting moderation, Feb 12, 2025)

## ACKNOWLEDGMENTS

We thank Emiliano Zamponi from the Polleux lab for helping with image annotation and the Polleux lab members for extensive discussions and annotations to generate the multi-user metrics. We thank Qiaolian Lu for her help with mouse colony management. Imaging was performed in the Zuckerman Institute’s Cellular Imaging platform. This work was supported by internal funding from the Zuckerman Institute, the NIH (5U19NS104649 and R35 NS127232), and a NOMIS Foundation Research Project.

## AUTHOR CONTRIBUTIONS

Conceptualization, S.B.G, F.P, L.A.H,; methodology development, S.B.G, A.P.S, L.A.H; data acquisition, annotation, and analysis, S.B.G, A.P.S, D.P., A.J.P, L.A.H, Investigation, S.B.G, F.P, L.A.H; writing— original draft, S.B.G, L.A.H; writing—review & editing, S.B.G, F.P, L.A.H; funding acquisition, F.P.; resources, F.P., L.A.H; supervision, F.P., L.A.H.

## DECLARATION OF INTERESTS

The authors declare no competing interests. geodesic distance computed along the dendrite in 3D (blue to white, proximal to distal). These validation outputs allow users to readily confirm accurate segmentation of dendritic spines and the parent dendrite. Scale bar, 5 µm.

## TABLES AND TEXT BOXES

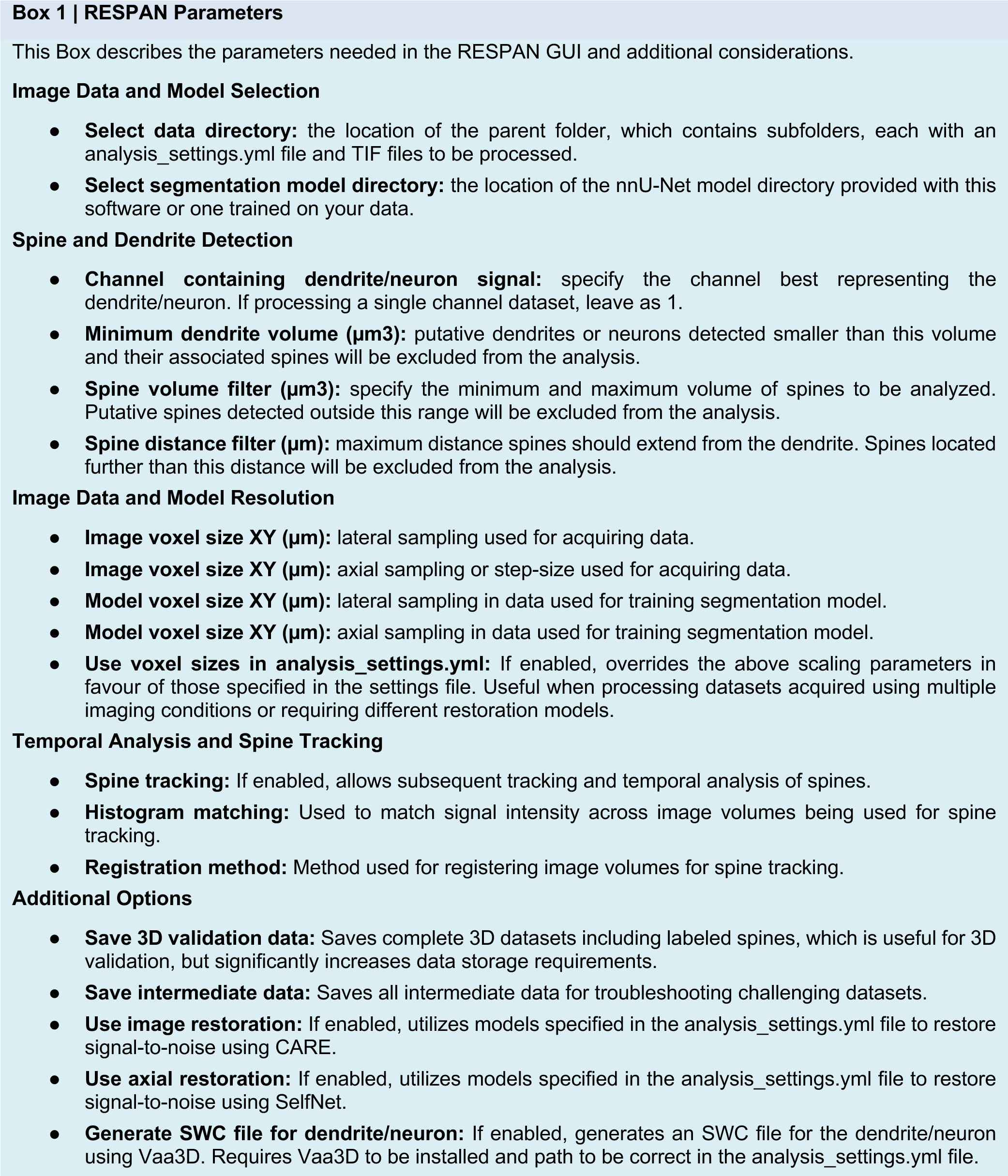

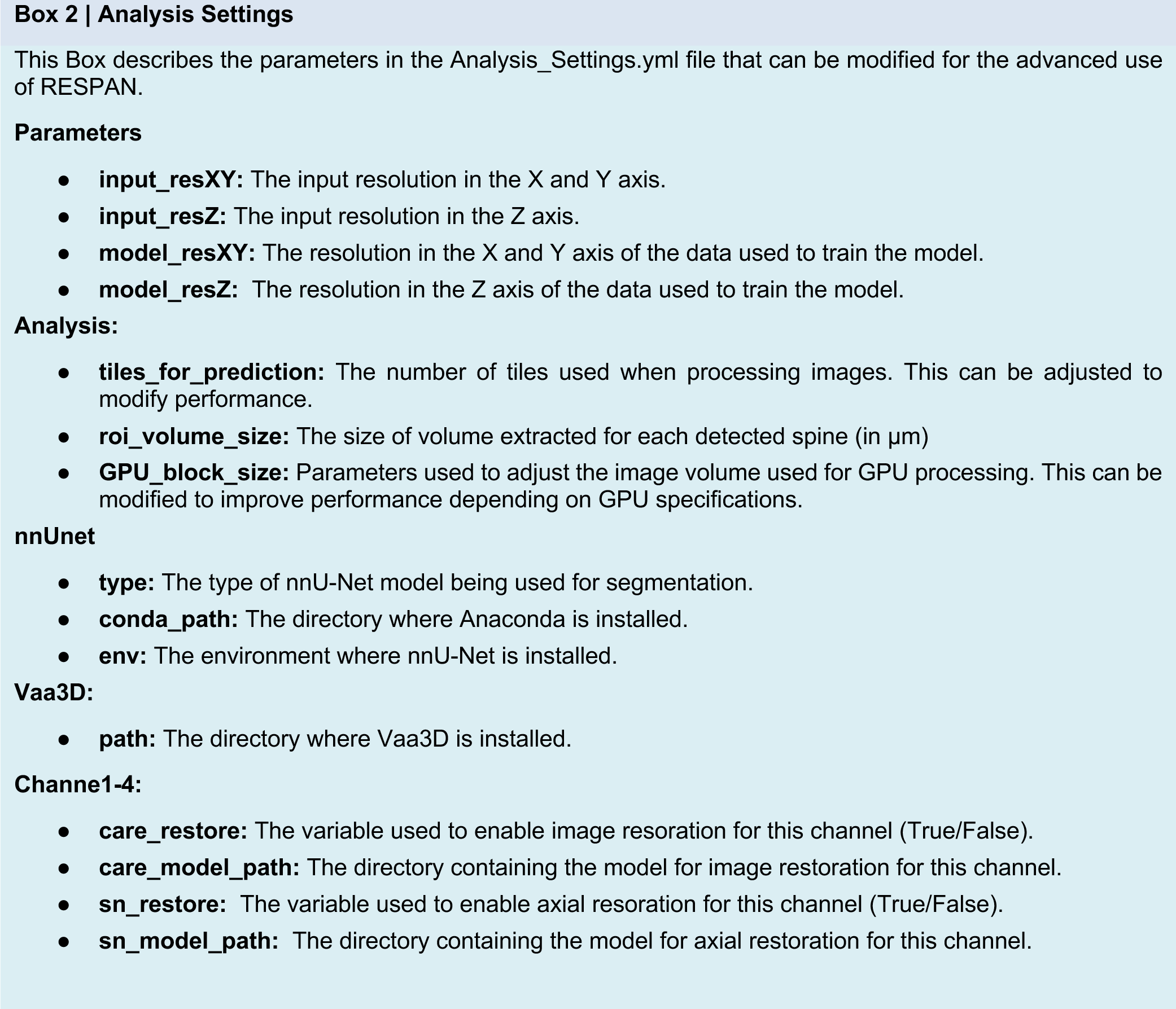

## METHODS

### KEY RESOURCES TABLE

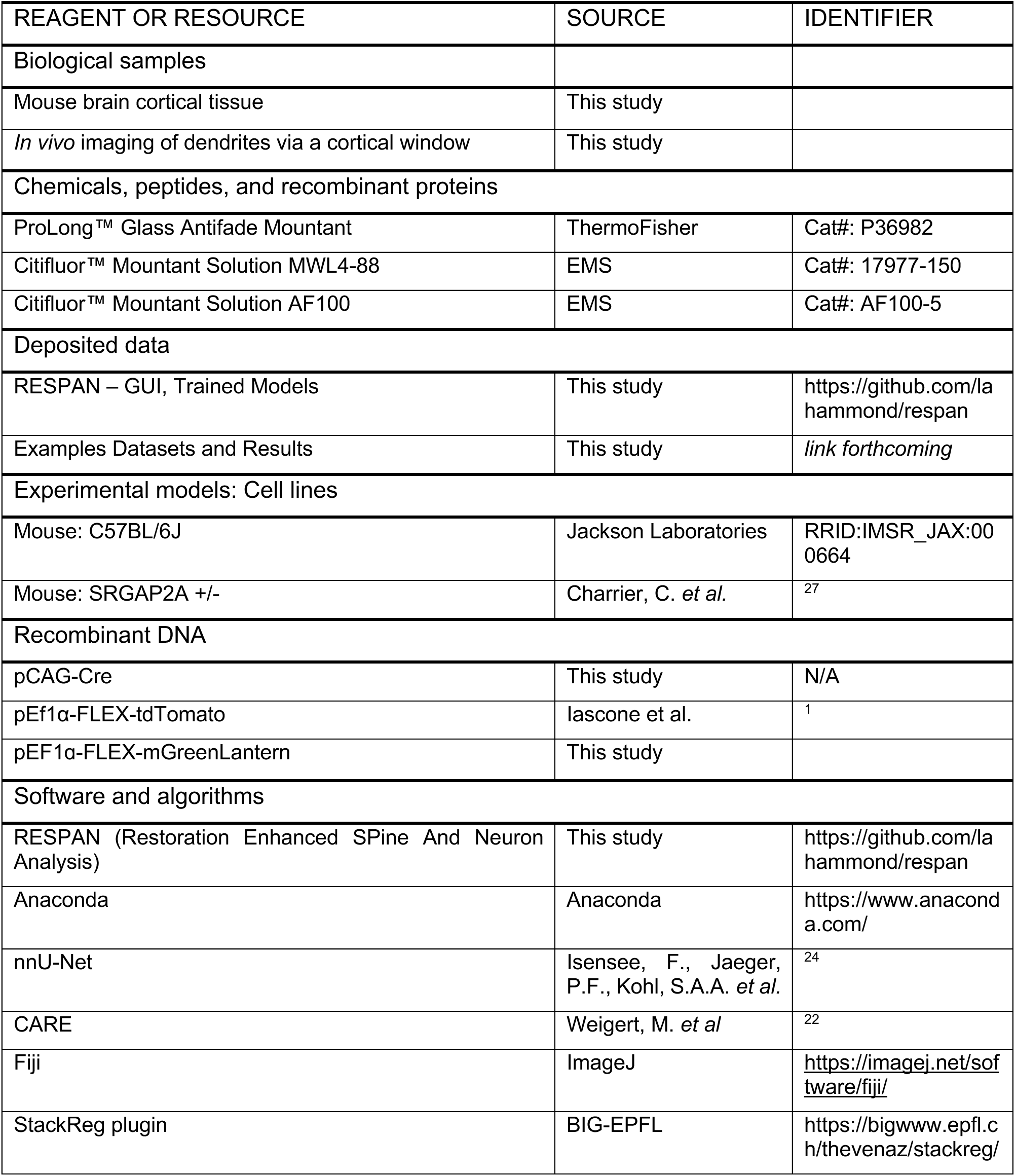
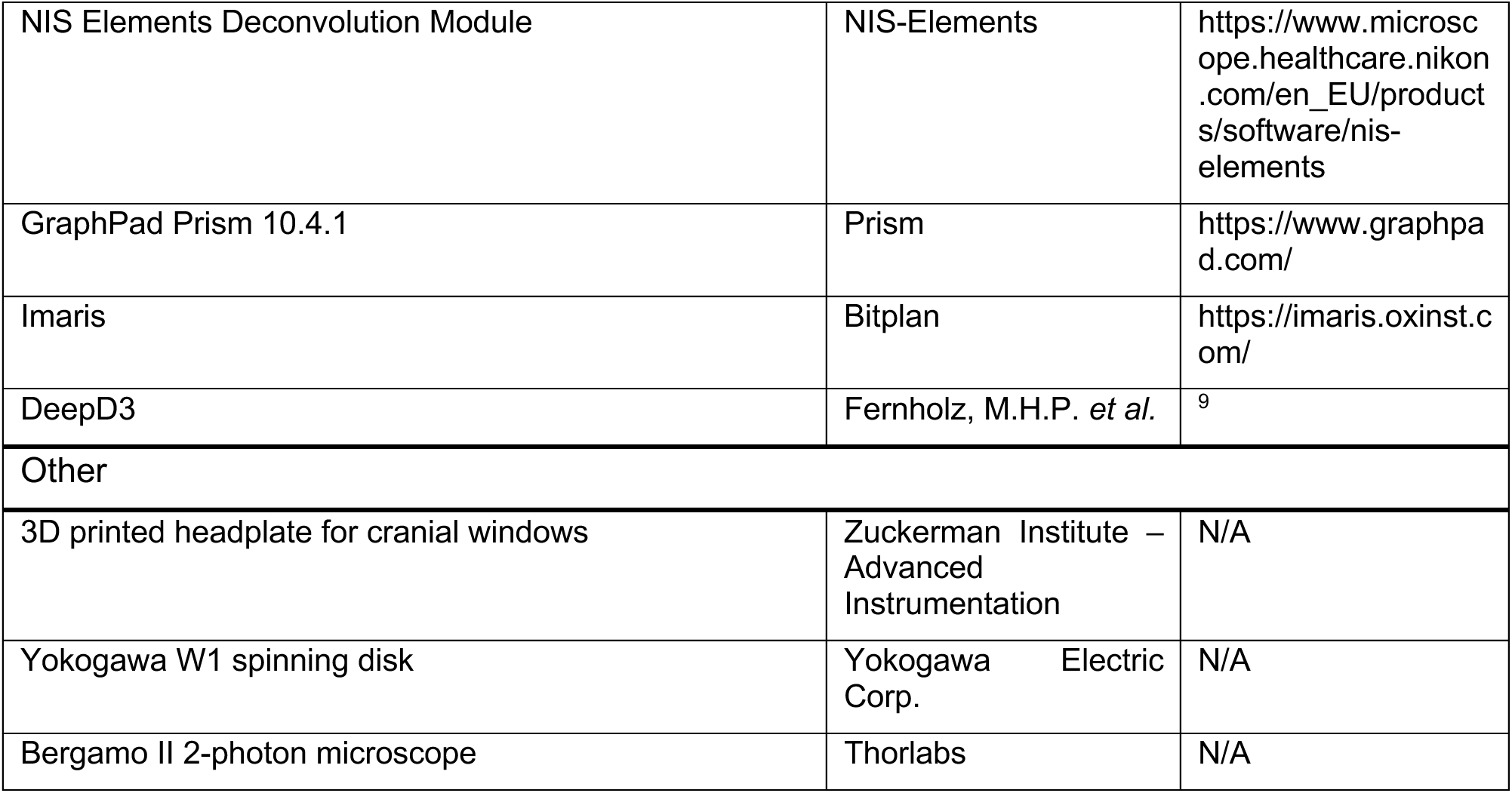

### EXPERIMENTAL MODEL AND STUDY PARTICIPANT DETAILS

#### Animals and Experimental Design

All animal procedures were approved by the Institutional Animal Care and Use Committee (IACUC) at Columbia University and conducted in accordance with institutional and federal animal welfare guidelines (IACUC Protocols AC-AABL2557 and AC-AABV8659). Mice were housed in a temperature-controlled environment on a 12-hour light/dark cycle, with ad libitum access to food and water.

#### In Vivo Imaging

Species/Strain: C57BL/6J (B6) and constitutive heterozygous knockouts for SRGAP2A+/- Genotype: WT (B6) and SRGAP2A+/-

Age/Developmental Stage: Mice were implanted with cranial windows at postnatal day (P)90 and imaged between P100–P120

Sex: Both male and female mice were used; sex was recorded for all animals

Housing and Maintenance: Standard housing conditions; individually housed post-surgery to prevent injury

Ethical Oversight: All procedures were approved by Columbia University IACUC

***Ex Vivo Imaging*** Species/Strain: C57BL/6J (B6) Genotype: WT

Age/Developmental Stage: Neurons were labeled in utero at embryonic day (E)15.5, with brains harvested at postnatal stages

Sex: Both male and female embryos were used; sex was not a variable in the analysis

Housing and Maintenance: Pregnant dams were housed under standard conditions until embryos reached E15.5 for in utero electroporation

Ethical Oversight: Procedures were approved by Columbia University IACUC

#### Sex as a Biological Variable

Both male and female mice were used in this study. While sex was recorded for all animals, the data were analyzed without sex-based segregation, as prior research indicates no significant sex-dependent differences in the measured parameters. However, potential sex-related effects are acknowledged as a limitation of this study.

### METHOD DETAILS

#### Animal Handling and Surgical Procedures

For in vivo imaging, mice underwent cranial window implantation at P90 under isoflurane anesthesia. A 4 mm circular craniotomy was performed over primary somatosensory cortex (S1), and a glass coverslip (No. 1, 4 mm diameter) was affixed with dental cement. A custom 3D printed headplate was attached to facilitate head fixation during imaging. Postoperative analgesia included subcutaneous injections of meloxicam (5 mg/kg). Mice were allowed 7–14 days for recovery before imaging.

For ex vivo experiments, in utero electroporation was performed at E15.5 to label Layer 2/3 pyramidal cortical neurons with fluorescent plasmids. Pregnant dams were anesthetized with isoflurane (1.5–2%), and embryos were injected with a plasmid mixture containing pCAG-Cre (1-10 ng/µL) and pCAG-FLEX- TdTomato (1 µg/µL). Plasmids were delivered into the lateral ventricle, followed by electroporation using five 50 ms pulses at 35 V (BTX ECM830 electroporator, 1 Hz frequency). Pups were allowed to develop postnatally before brains were extracted for imaging.

#### Tissue Preparation for Ex Vivo Imaging

Mice were anesthetized with isoflurane and transcardially perfused with ice-cold phosphate-buffered saline (PBS) followed by 4% paraformaldehyde (PFA) in PBS. Brains were dissected and post-fixed in 4% PFA overnight at 4°C, then rinsed in PBS and sectioned at 100 µm using a Leica VT1000 S vibratome. Sections were mounted in ProLong Gold antifade reagent for imaging.

#### In Vivo Two-Photon Imaging

Imaging was performed using a Bergamo II two-photon microscope (Thorlabs) equipped with a Ti:Sapphire laser (Coherent Chameleon Ultra II, 920 nm excitation). A 25× 1.0 NA water-immersion objective (Olympus, XLPlan N) was used for data acquisition. Image stacks were acquired at 1024 × 512 pixels with a pixel size of 0.102 µm and 1 µm axial step size. The laser operated at an 80 MHz repetition rate with a 100 fs pulse width, and power at the sample was maintained at <15 mW to minimize phototoxicity. Mice were head-fixed on a custom stage, and imaging sessions lasted <60 min per session to prevent physiological stress, minimize photodamage, and minimize exposure to anesthetic.

#### Spinning Disk Confocal Imaging (Ex Vivo Samples)

Sections were imaged using a Yokogawa W1 spinning disk confocal system mounted on a Nikon Ti2 inverted microscope controlled with NIS-Elements. Images were acquired with a 100× 1.35 NA SR HP Plan Apo silicone immersion objective and a Prime BSI sCMOS camera (Teledyne Photometerics), resulting in pixel size of 65nm and a z-step size of 150nm. 488 and 561 nm laser lines were used to sequentially excite GFP and TdTomato, respectively, with matched bandpass emission filters installed in a triggered high-speed filter wheel (IDEX Health & Science LLC). When appropriate, images were further processed using a blind deconvolution algorithm with 25 iterations using the low noise parameter in the NIS Elements.

#### Data Acquisition and Processing

In vivo time-lapse imaging was performed via daily imaging over 2 continuous days to track dendritic spine dynamics across a 24 hour time interval. Ex vivo datasets were processed for 3D reconstruction and dendritic spine analysis using RESPAN (Restoration Enhanced SPine and Neuron Analysis), which integrates content-aware image restoration with deep learning-based segmentation. Motion correction for in vivo imaging was conducted using StackReg (https://github.com/fiji-BIG/StackReg) to compensate for minor movement artifacts. Spine detection and morphometric analyses were performed using nnUNet-based segmentation models, which were trained on a diverse dataset of manually annotated spines to ensure robustness across varying imaging conditions. Quantifications included spine density, spine head volume, and spine-to-dendrite distance, with statistical validation performed against expert-annotated ground truth datasets.

### QUANTIFICATION AND STATISTICAL ANALYSIS

All statistical analyses were performed using GraphPad Prism 9.0. Statistical details for each experiment, including sample sizes (n), measures of central tendency, variability, and statistical significance, are reported in the figure legends and corresponding text in the Results section.

#### Statistical Tests and Data Distribution

To compare differences in dendritic spine density, spine length, and volume between WT and SRGAP2A+/- mice, we used Kolmogorov-Smirnov tests, a non-parametric method suitable for comparing distributions without assuming normality. For comparisons of imaging-based segmentation performance (e.g., spine detection accuracy, precision, recall, and F1 score), we employed paired two- tailed t-tests and Wilcoxon matched-pairs signed-rank tests, as appropriate. For datasets involving multiple comparisons, two-way ANOVA followed by Benjamini-Krieger-Yekutieli correction was applied to control the false discovery rate (FDR).

#### Definition of Sample Size (n)

For *in vivo* imaging, each n represents an individual dendritic segment imaged from a single neuron, and multiple neurons were imaged per animal. For ex vivo experiments, each n corresponds to an apical dendritic segment from a single neuron; multiple neurons were imaged from multiple mice. Final sample sizes were based on prior studies assessing dendritic spine morphology in similar experimental paradigms and are reported in figure legends.

#### Randomization and Blinding

Mice were randomly assigned to experimental groups at the time of imaging, and all spine analyses were performed blinded to genotype to reduce bias. Image processing and quantification via RESPAN also remained blind to the genotype to minimize manual intervention and ensure unbiased analysis.

#### Segmentation Validation and Accuracy Metrics

RESPAN’s automated segmentation performance was validated against ground truth expert annotations, and performance was quantified using Dice coefficient, F1 score, precision-recall curves, and Hausdorff distance. Object-level segmentation accuracy was assessed by comparing IoU (Intersection over Union) values with a threshold of 0.50 to define correctly detected spines.

#### Statistical Significance Criteria

For all analyses, statistical significance was defined as p < 0.05, with exact p-values reported in the figures and results section. Data distributions were examined for normality, and appropriate parametric or non-parametric tests were applied based on distribution characteristics.

### ADDITIONAL RESOURCES

RESPAN source code and application: https://github.com/lahammond/respan

## SUPPLEMENTAL INFORMATION

**Supplemental Figure 1.**
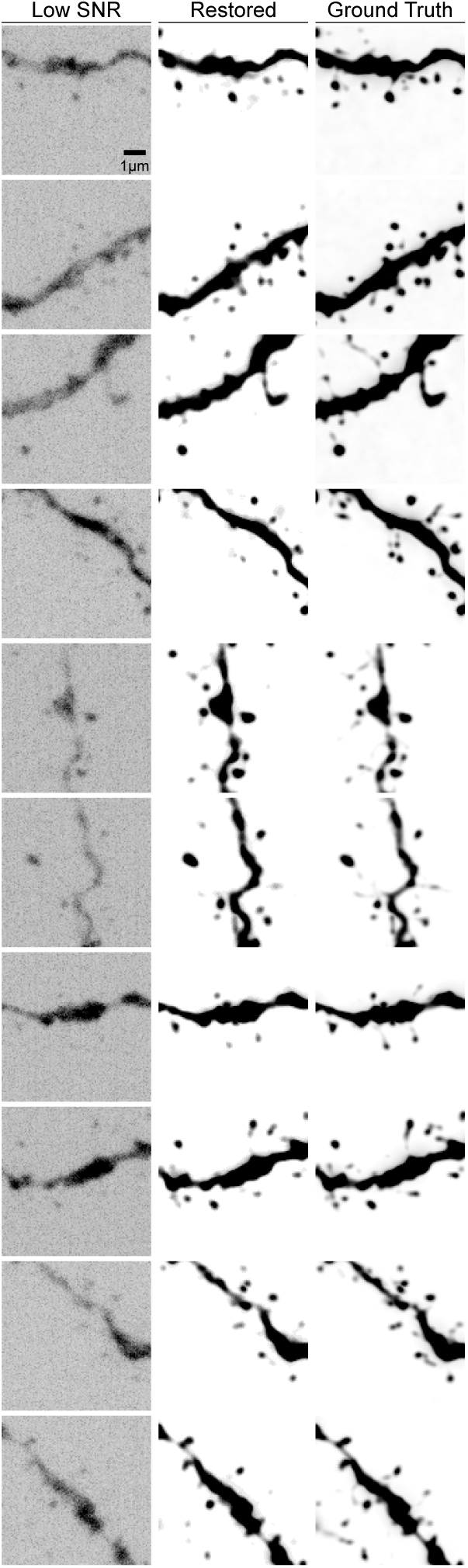
R**e**presentative **Examples of CARE Restoration** Isolated maximum intensity projection views of a dendritic segment are shown under three conditions: low-SNR acquisition (left), CARE-restored output (middle), and high-SNR ground truth data (right). Each row demonstrates content-aware restoration of signal and contrast, revealing spine heads, necks and fine dendritic features that are difficult to discern in low-SNR images. Scale bar, 1 µm.

**Supplemental Figure 2.**
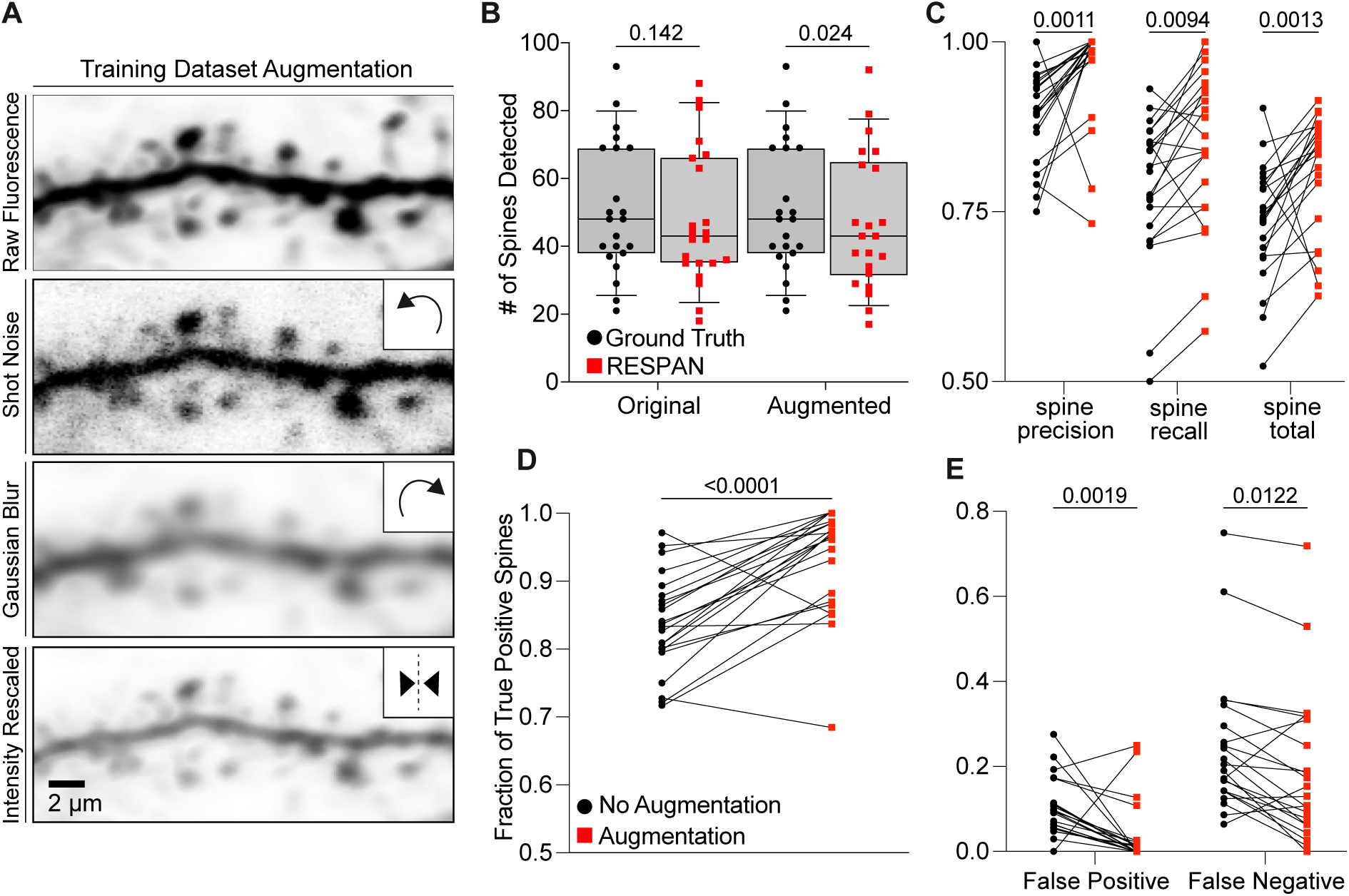
*Enhancement of Spine Detection Performance Through Data Augmentation*. (A) Data augmentation techniques applied during model training, illustrated through representative images: original raw image and subsequent transformations including shot noise, Gaussian blur, and pixel intensity scaling, aimed at enhancing model robustness and generalizability; random rotations and flips were performed to the data to avoid overfitting during model training. Scale bar, 2 µm. (B) Comparison of the number of spines detected in original and augmented datasets by ground truth (black) and RESPAN (red), analyzed using two-way ANOVA, indicating no significant difference (ns) to significant differences (*p < 0.05, ****p < 0.0001) in detection counts.(C) Scatter plots comparing spine detection metrics—precision, recall, and total performance—between original and augmented datasets, analyzed using two-way ANOVA with statistical significance indicated (**p < 0.01). (D) Bar graph showing the fraction of true positive spines detected, comparing models trained with no augmentation (black) and with augmentation (red), analyzed using the Wilcoxon matched-pairs signed rank test. The asterisks indicate a statistically significant increase in the detection of true positives (****p < 0.0001). (E) Scatter plot illustrating the reduction in false positives and false negatives in augmented datasets, analyzed using two-way ANOVA, with statistical tests indicating significant differences in error rates (**p < 0.01, *p < 0.05).

**Supplemental Figure 3.**
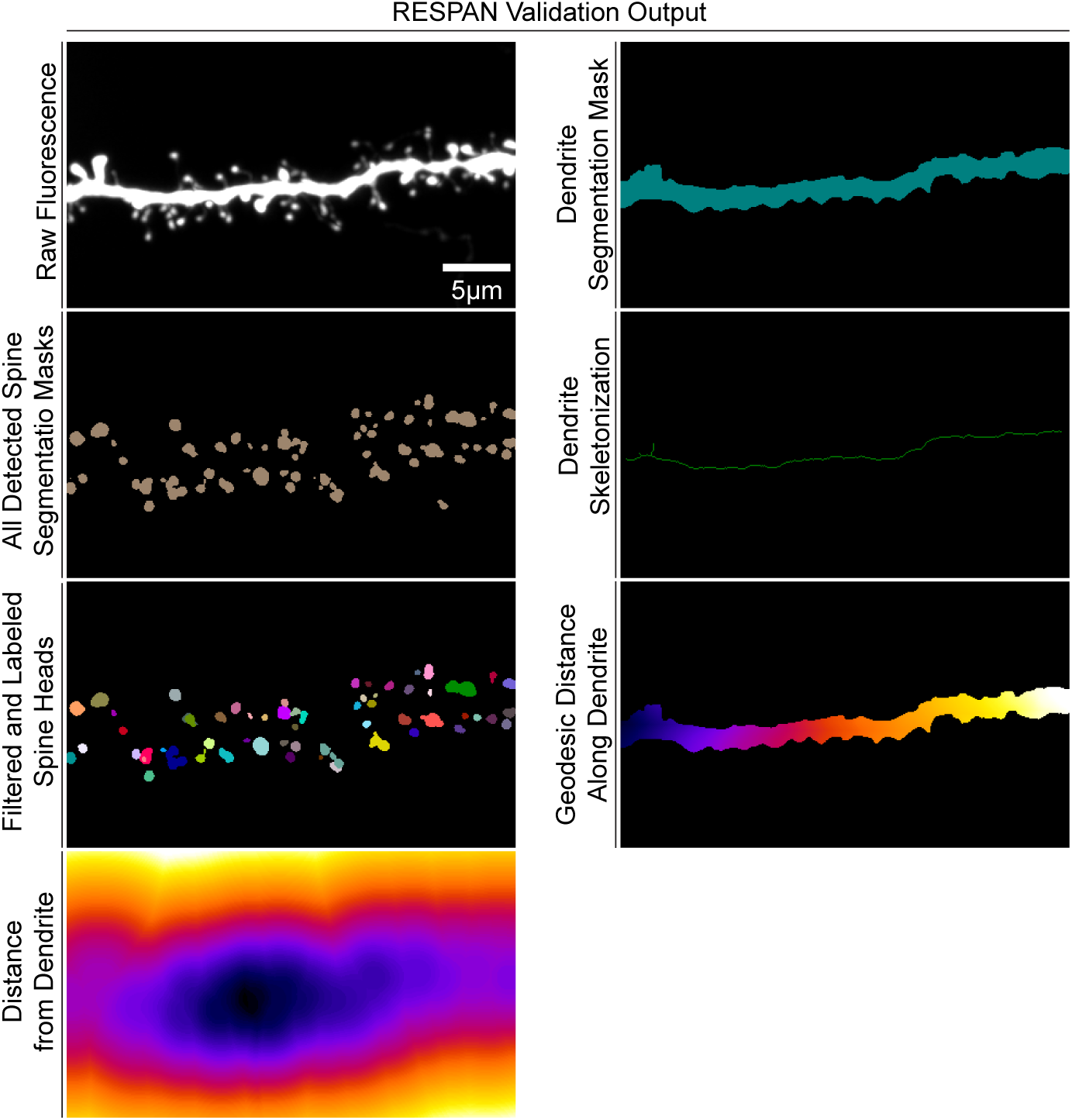
R**E**SPAN **Validation Output for Dendritic Spine Segmentation** Maximum intensity projection views generated for validating RESPAN outputs. (Left) From top to bottom: raw fluorescence image of the analyzed dendritic segment; all detected spine segmentation masks; color-coded filtered spine segmentation masks after applying morphological criteria (minimum/maximum volume; maximum distance from dendrite); and a heatmap illustrating each voxel’s distance from the dendrite shaft (cooler hues, closer distance; warmer hues, greater distance). (Right) Corresponding dendrite segmentation mask, dendrite skeletonization, and color-coded

## Notes

### Competing Interest Statement

The authors have declared no competing interest.

### Summary of Updates

We improved the GUI/computer interface for this machine-learning based approach. We also improved some of the output analysis performed using RESPAN including a revised Figure 2, a new Figure 3 and a new Figure 6. We also provide more details on how to use the GUI interface.

